# CDRxAbs: Antibody Small-Molecule Conjugates with Computationally Designed Target-Binding Synergy

**DOI:** 10.1101/2023.12.19.572259

**Authors:** Jingzhou Wang, Aiden J. Aceves, Stephen L. Mayo

## Abstract

Antibody-drug conjugates (ADCs) combine the advantages and offset the disadvantages of their constituent parts to achieve a refined spectrum of action. We combine the concept of ADCs with the full atomic simulation capability of computational protein design to define a new class of molecular recognition agents: CDR-extended antibodies, abbreviated as CDRxAbs. A CDRxAb incorporates a covalently attached small molecule into an antibody/target binding interface using computational protein design to create an antibody small-molecule conjugate that binds tighter to the target of the small molecule than the small molecule would alone. CDRxAbs are also expected to increase the target binding specificity of their associated small molecules. In a proof-of-concept study using monomeric streptavidin/biotin pairs at either a nanomolar or micromolar-level initial affinity, we designed nanobody-biotin conjugates that exhibited >20-fold affinity improvement against their protein targets with step-wise optimization of binding kinetics and overall protein stability. The workflow explored through this process promises a novel approach to optimize small-molecule based therapeutics and to explore new chemical and target space for molecular-recognition agents in general.

**Significance:** We defined a general method for optimizing molecular recognition reagents that involve small molecules and demonstrated an application of this method using a model system. Instead of using traditional approaches for modifying a small molecule to improve its binding properties, we use computational protein design to build an antibody/small molecule conjugate that allows the target-binding strength (and specificity) of the small molecule to be tuned through changes in the amino acid sequence of the antibody scaffold. This method introduces a novel approach for optimizing the binding properties of small molecules and expands the potential application scenarios for antibody-drug conjugates.

## Introduction

The mechanisms of action of most pharmaceuticals involve drug-target interactions that are mediated by synthetic small molecules or monoclonal antibodies [1,2]. Despite impressive successes, many biological pathways are still considered to be “undruggable” because the difficulty of engineering the desired interactions or the pharmacological trade-offs for establishing the interactions outweigh the potential benefits [3–6]. This observation drives creation of new approaches and modalities for addressing challenging targets, which can also expand the targetable molecular space itself [7]. To create new drug modalities, a common approach is to combine existing modalities in order to capitalize on individual advantages and offset individual flaws [7,8]. Antibody-drug conjugates (ADCs), for example, take advantage of the high specificity and biological compatibility of monoclonal antibodies to improve therapeutic indices of small-molecule drugs [8,9]. The antibody and small-molecule drug components of ADCs are typically developed separately and bind to different targets during their course of action [8,9]. Generally, ADCs improve the specificity of their conjugated drugs by delivering the small molecule payloads into target cells through specific antibody-induced receptor endocytosis [9,10]. Some ADCs and peptide-drug conjugates have also been reported to improve the metabolic stability, circulation half-life, and solubility of linked small molecules through antibody-associated pharmacokinetics, chemical environment around the conjugation sites, and linker design, indicating that protein conjugation could modulate a wide range of small-molecule properties [10–14].

Recently, Cheng *et al.* developed ADCs whose antibody and drug components bind to an identical protein target to achieve synergistic binding/inhibition effects [15]. In their study, the co-crystal structure of a small molecule drug, sitagliptin, a separately-developed antibody, 11A19, and the protein target, DPP- IV, was solved first [16]. Based on the structure, optimal conjugation sites and linker sequences were then searched to create ADCs that exhibited 13 to 32-fold IC50 improvement relative to sitagliptin alone against the target [15]. Cheng *et al.*’s work demonstrated that small molecule binding could be improved through conjugating to antibodies, suggesting the potential to expand the chemical space and therefore target space of molecular recognition agents that involve small molecules. However, the approach requires the independent development and characterization of the individual antibody and small molecule components and doesn’t incorporate small-molecule binding directly into the antibody/antigen interface, which could be advantageous for certain applications. In order to realize the full potential of this approach, a workflow that can efficiently identify a compatible antibody sequence and conjugation strategy for a desired small molecule binding event would be ideal.

Recently, Lewis *et al.* explored a similar concept of fibronectin small-molecule conjugates, where the fibronectin component was engineered by yeast display in presence of the bioconjugation [17]. The engineered conjugates displayed 9 and 80-fold improvement respectively in potency and selectivity against enzyme targets, further strengthening the hypothesis that a small molecule binding behavior could be directly tuned by a conjugated protein sequence.

In this study, we explored the feasibility of computationally designing the antibody and linker components of synergistically-binding ADCs. We introduced the concept of CDR-extended antibodies (CDRxAbs), which are antibodies whose complementary-determining regions (CDRs) contain a covalently attached small molecule ligand that binds to a desired target with surrounding CDR amino acid sequences tailored to strengthen the target-binding interactions (Figure 1A). For this proof-of-concept study, we focused our design on nanobodies, which are llama-derived single-domain antibody fragments [18–20]. Using a modified streptavidin-biotin interaction pair as model system, we demonstrated that with only the structural knowledge of small-molecule/target interactions, nanobody small-molecule conjugates can be computationally designed to bind a target tighter than the small molecule itself. Through subsequent computationally-directed sequence design, the affinity, binding kinetics, and overall stability of the conjugates can be improved in a step-wise manner. Greater than 20-fold affinity improvements together with targeted kinetic-tuning were achieved when the starting small-molecule/target affinity is as weak as 1 µM or as strong as 7 nM. Exploration of various computational methods revealed key design principles from which we proposed a general design strategy for this novel molecular modality.

**Figure 1:**
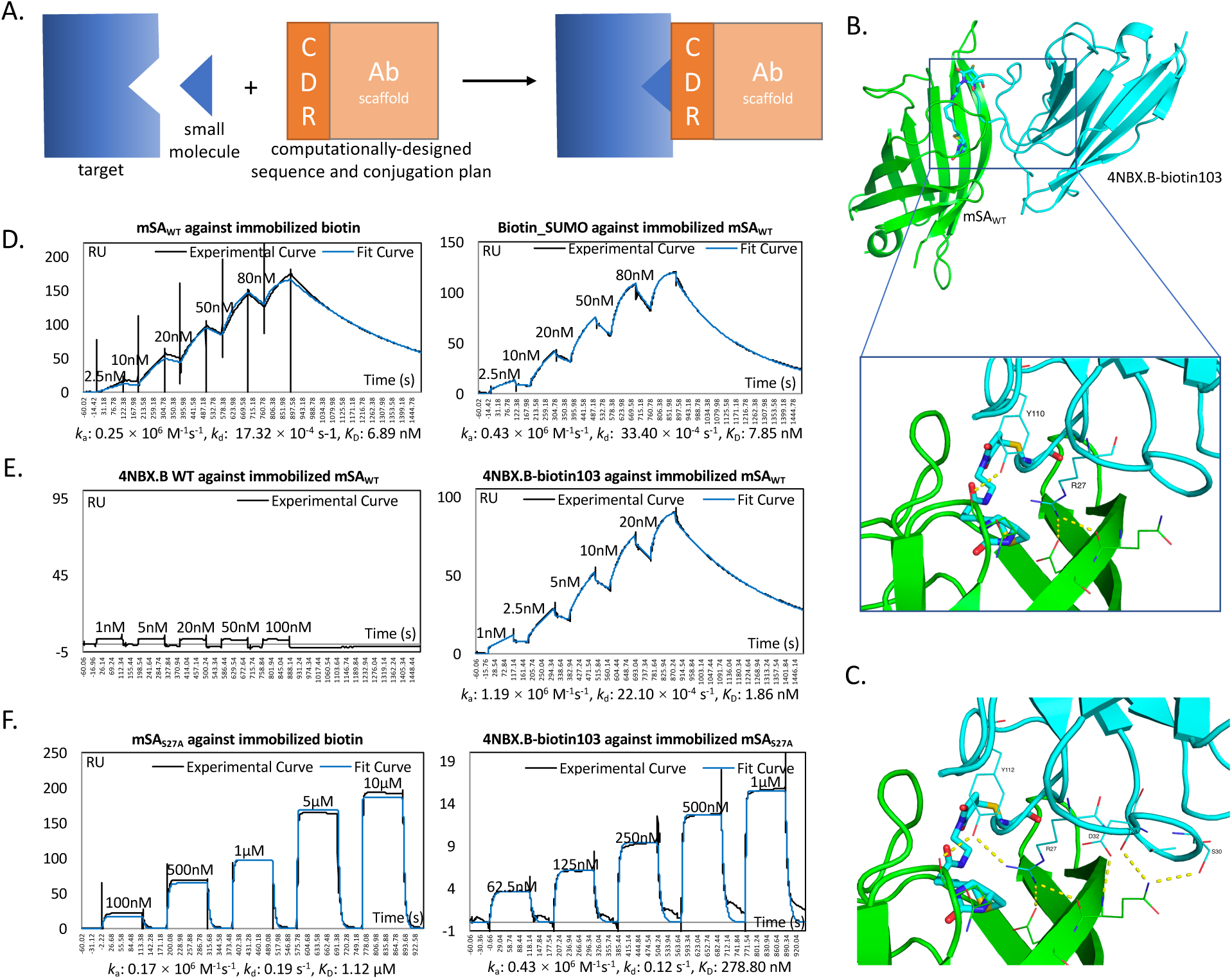
Computationally-designed nanobody-biotin conjugates. (A) Schematic representation of workflow. (B) Final model of 4NBX.B-biotin103 in complex with mSA_WT_ streptavidin. mSA_WT_ shown in green; nanobody shown in cyan; Biotin103 side chain shown as stick. (C) Potential hydrogen bond interactions between nanobody and mSA_WT_. (D) SPR measurement of mSA_WT_/biotin binding with immobilized biotin (left) and immobilized mSA_WT_. (E) SPR measurement of 4NBX.B-WT/mSA_WT_ binding (left) and 4NBX.B-biotin103/mSA_WT_ binding (right). (F) SPR measurement of mSA_S72A_/biotin (left) and 4NBX.B-biotin103/mSA_S72A_ binding (right).

## Results and Discussion

### Computationally-designed nanobody small-molecule conjugation creates tighter binders against the small-molecule target protein

We first asked whether computationally-determined nanobody sequences and their designed conjugation to a small molecule can exhibit an enhanced binding affinity to the small-molecule target. Designing the antibody components of synergistically-binding ADCs involves creating a new antibody/target interface, which is challenging because of the difficulty in predicting the conformation of antibody CDR loops when bound to a target protein and the general challenge of accurately modeling long structured loops [21–24]. To avoid the challenge of accurately modeling CDR conformations, we adopted an approach similar to anchored-design methods [25]. Anchored-design creates new protein-protein interfaces by first identifying hotspot residues that favorably interact with the target, then designing protein scaffolds to stabilize the anchoring hotspots [25–27]. For synergistically-binding ADCs, the conjugated small molecule is defined as a hotspot “residue” that interacts with the target protein. Therefore, to create co-targeting ADCs, the small molecule can be designed as an anchoring noncanonical CDR residue with binding interactions strengthened by additional CDR/target interactions, integrating the drug/target interaction into the antibody/target binding event and biasing the CDRs to adopt the designed conformation.

We divided the design strategy into the following two steps: predicting the optimal CDR binding poses against the target in the small molecule binding region and searching for a conjugation strategy that accommodates both the optimized CDR pose and the target small molecule interaction. As a proof-of-concept we chose monomeric streptavidin as a target and biotin as the small molecule. Streptavidin/biotin interactions have been structurally well characterized. Moreover, monomeric streptavidin variants have been reported with biotin binding affinities reduced by more than 10^5^-fold relative to the tetrameric streptavidin parent [27,28] which allows for a range of initial starting affinities.

To search optimal CDR binding conformations, we first docked various nanobody scaffolds onto a monomeric streptavidin structure with a computationally-modeled triple mutant, S45A/T90A/D180A, that was reported to monomerize streptavidin and reduce the biotin-binding affinity to 1.7 µM [28]. We then performed loop-modeling on docked poses in an attempt to optimize CDR conformations against the target surface. Most of the top loop modeling solutions were not representative of naturally occurring interactions (data not shown). In order to constrain the search to sample viable CDR structures, we restricted the search to previously observed nanobody CDR binding conformations [30]. We curated a library of 154 nanobody structures with diverse target-binding CDR conformations from the Protein Data Bank (PDB) and individually docked them onto the target streptavidin surface (Fig. S1A). 2310 docked poses were generated and filtered to identify the most realizable binding conformations, resulting in 7 final binding poses (Fig. S1C). Optimal conjugation strategies were then searched on the finalized poses. We chose to conjugate biotin to nanobody CDRs via cysteine-maleimide chemistry, a commonly used conjugation method in ADCs (Fig. S2A) [31]. Biotin C2 maleimide was selected as the conjugation reagent. Optimal nanobody scaffolds and conjugation sites were determined by computationally screening a rotamer library of the cysteine-conjugated side chain at various CDR positions on the nanobody/streptavidin poses. The top-ranked conjugation plan was amino acid position 103 of the 4NBX.B nanobody scaffold (chain B of PDB structure 4NBX), which natively binds to the bacterial toxin TcdA, a target unrelated to streptavidin [32]; the resulting molecule is named 4NBX.B-biotin103. The computationally relaxed complex shows hydrogen bonds between the sidechains of residues Y112 and R27 on 4NBX.B-biotin103 and the backbone of streptavidin. In the native 4NBX.B complex with TcdA, these two residues also participate in intermolecular sidechain/backbone hydrogen bonds, suggesting that the designed pose is closely related to the natural binding mode of 4NBX.B and that these interactions could potentially stabilize the biotin-modified nanobody’s interactions with streptavidin. (Fig. S2B-C).

We then synthesized 4NBX.B with site 103 mutated to cysteine, and coupled it with biotin C2 maleimide. We attempted to purify and refold the S45A/T90A/D128A mutant of streptavidin to perform binding measurements, but the molecule was insoluble in our hands (data not shown). Therefore, we structurally aligned our triple-mutant streptavidin model and a different previously reported monomeric streptavidin variant, mSA (57% sequence pairwise identity) [29]. The C-alpha atom RMSD for the alignment was 0.5Å. We then relaxed the resulting complex of 4NBX.B-biotin103 and mSA (Fig. S2D). The 4NBX.B-biotin103/mSA model preserved the rotamer configuration of the conjugated biotin from the triple-mutant streptavidin model as well as the intermolecular Y112 and R27 hydrogen bonds (Fig. S2D).

Surface plasmon resonance (SPR) binding experiments conducted at 25 ℃ confirmed that 4NBX.B-biotin103 binds to immobilized mSA with a *K*_D_ of 1.8±0.1 nM compared to mSA binding to immobilized biotin with a *K*_D_ of 7.0±0.1 nM, indicating a moderate 4-fold affinity improvement that is largely due to a higher *k*_a_ (Fig. 1D-E, 5A). Wildtype 4NBX.B did not show binding to immobilized mSA at concentrations up to 100 nM, indicating that the biotin in the 4NBX.B-biotin103 conjugate is required for binding to mSA (Fig. 1E left panel). The SPR-measured biotin/mSA affinity is similar to previously-published fluorescence polarization spectroscopy data: 2.8±0.5 nM at 4 ℃ and 5.5±0.2 nM at 37 ℃ [29]. However, given the lower data-fitting quality of the mSA/biotin binding curves relative to the 4NBX.B- biotin103 binding curves, we performed an alternative determination of the mSA/biotin affinity by binding Smt3 SUMO protein biotinylated at the N-terminus with biotin C2 maleimide to immobilized mSA. Smt3 SUMO protein has an unstructured N-terminus that we hypothesized would minimize the interaction between the protein components [33]. The data quality was improved and the resulting *K*_D_, 9.6±1.2 nM, is similar to what we determined above (Fig. 1D right panel).

To determine whether computationally-designed nanobody conjugation shows improved affinity when starting with weakly-binding small molecules, we created a single mutation of mSA, mSA_S27A_, whose counterpart S45A in wild type streptavidin reduces biotin-binding strength and was predicted by molecular dynamics (MD) simulation to minimally affect the overall structure [34]. Size-exclusion chromatography (SEC) shows that mSA_S27A_ elutes at the same time as mSA_WT_ (Fig. S3A). SPR measurements show that mSA_S27A_ binds to biotin with a *K*_D_ of 1.14±0.02 µM, while 4NBX.B-biotin103 binds to mSA_S27A_ with a *K*_D_ of 245±41 nM, indicating a similarly-moderate 5-fold improvement in binding affinity (Fig. 1F, 5B). Together, the above results demonstrate that a purely computational method based solely on structural information of the individual components can be used to design CDRxAbs that exhibit affinity-enhancing synergistic binding effects.

### Sequence design improves the binding affinity and kinetics for computationally designed conjugates

Next, we performed computational sequence design on the CDR loops of 4NBX.B-biotin103 with the goal of improving its binding affinity to mSA. We performed *in silico* site-saturation mutagenesis on each CDR position in the complex to identify positions tolerant to mutation, resulting in a set of 9 positions (SI Appendix). Computational sequence designs using combinations of four identified CDR positions each were tested in parallel. The amino acids allowed for each position in these calculations were set based on the known sequence diversity of nanobody CDR loops [35]. Analysis of the design output revealed that designs with positions 31, 32, 104, and 105 most frequently returned sequences that were predicted to form additional intermolecular hydrogen bonds and that were also energetically stable. The top-ranked variant by energy, v119, has CDR1 mutations M31H and D32A, and CDR3 mutations N104S and W105H. New hydrogen bonds were predicted between H31 and Q108 of mSA and between H105 and E105 of mSA (Table S1, Fig. 2A). The D32A mutation eliminates a buried, unpaired charged residue. SPR measurements show that the v119 variant binds mSA_WT_ with a *K*_D_ of 0.9±0.2 nM, indicating an ∼2-fold improvement from 4NBX.B-biotin103_WT_ (Fig. 2A, 5A). The *K*_D_ improvement was primarily due to an increase in *k*_a_ (Fig. 2A, 5A). With the goal of reducing *k*_d_, we selected variant v149, which has the largest number of predicted hydrogen bonds, 15, of the top 20 ranked sequences. The v149 variant has mutations M31R, D32S, N104A, and W105R that were predicted to form extensive interactions with Y96, E105, and Q108 of mSA, including a potential salt bridge between nanobody R105 and mSA E105 (Fig. 2B). Interestingly, R-E interactions appear to be frequently used by nanobodies [36]. Compared to v119, SPR measurements for v149 show an ∼2-fold lower *k*_d_ and an ∼4-fold higher *k*_a_ that together contribute to a *K*_D_ of 0.12±0.01 nM, a >20-fold *K*_D_ improvement compared to biotin/mSA_WT_ affinity (Fig. 2B, 4A). However, SEC analysis of v149 shows signs of aggregation (Fig. 3A).

**Figure 2:**
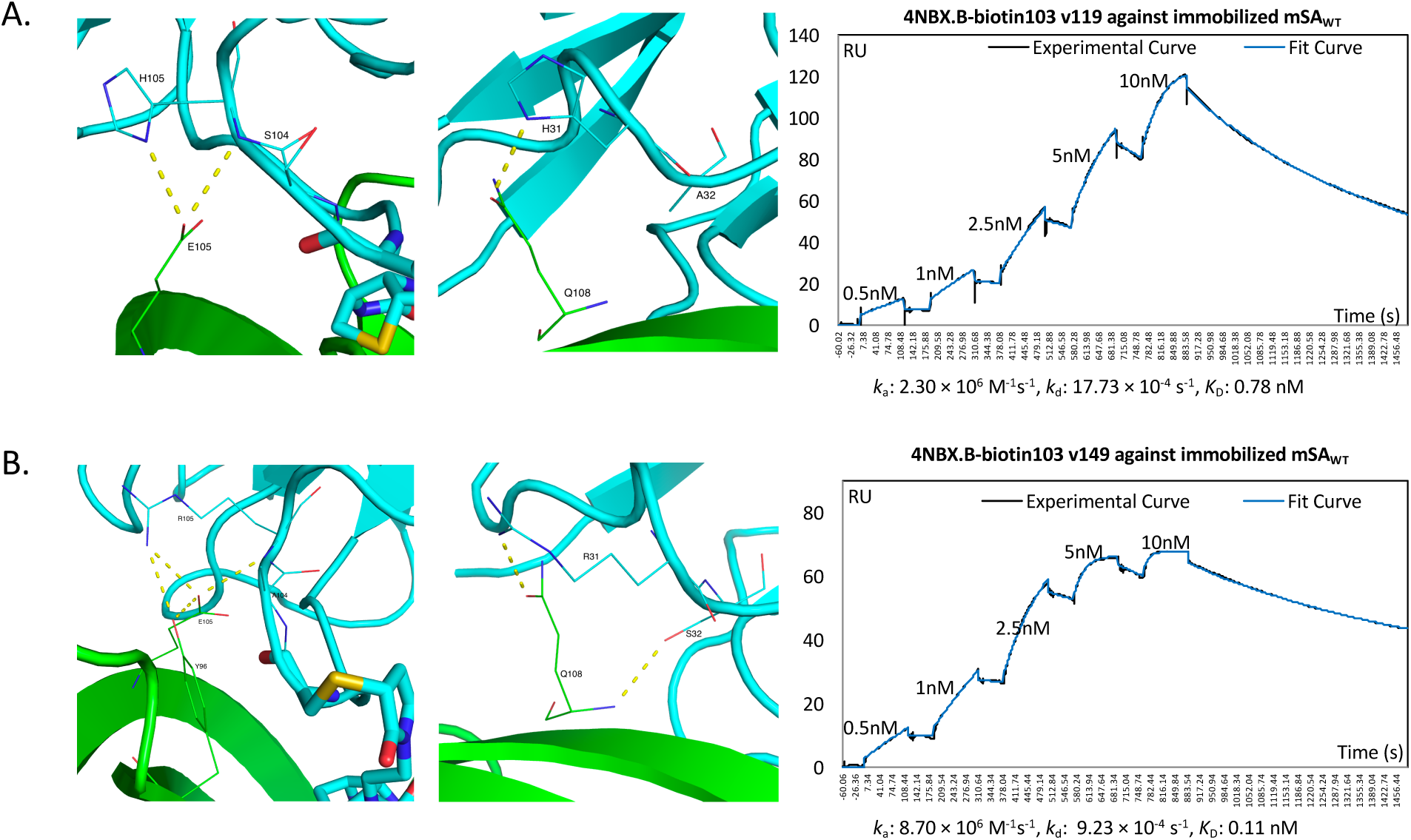
Design models and SPR binding for CDR sequence designs v119 and v149. (A) v119 design showing potential inter-protein hydrogen bonds (left and middle -- mSA_WT_, green; v119, cyan; Biotin103 thick sticks) and SPR measurement of v119/mSA_WT_ binding (right). (B) v149 design showing potential inter-protein hydrogen bonds (left and middle) and SPR measurement of v149/mSA_WT_ binding (right).

**Figure 3.**
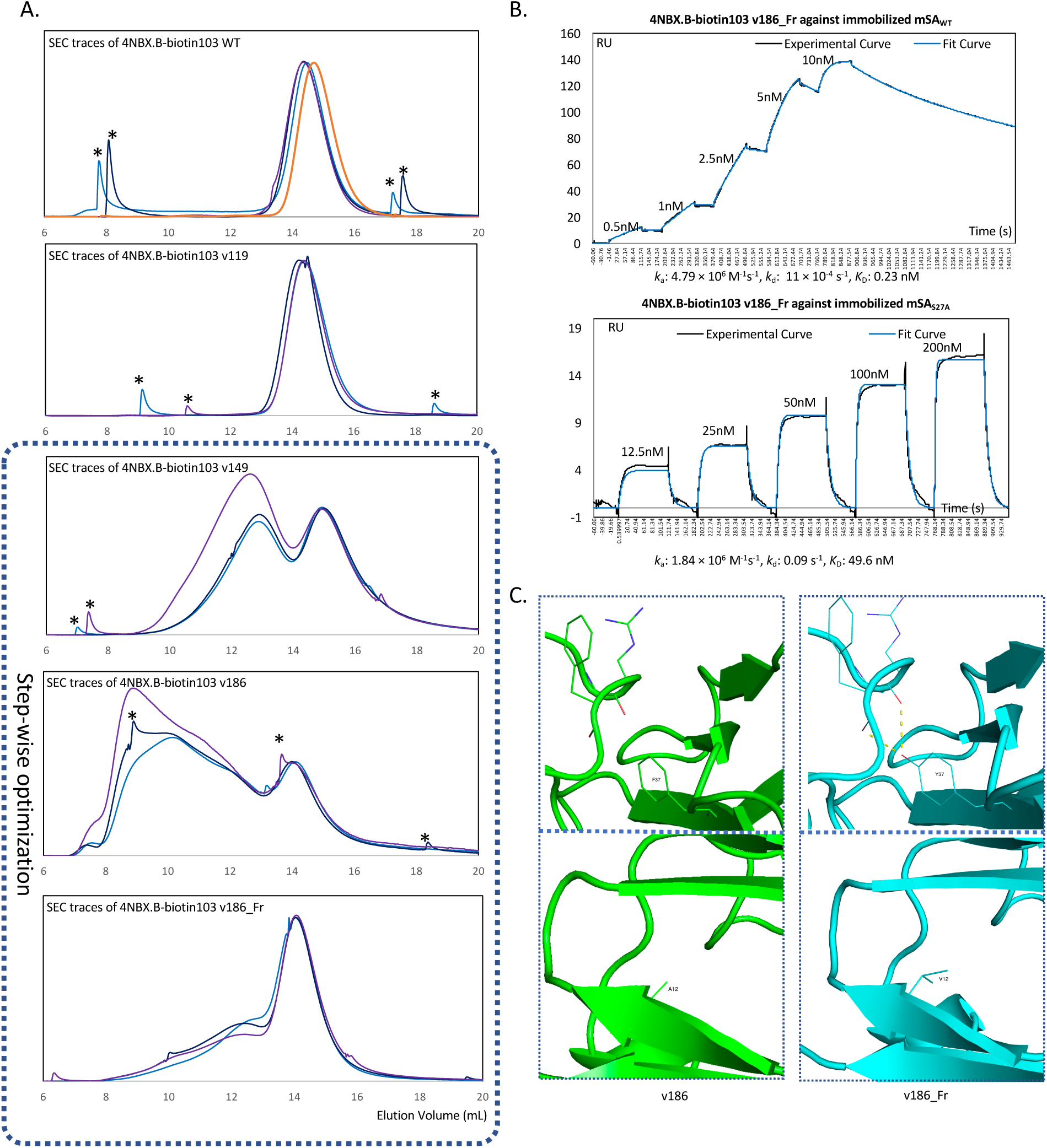
Framework design for reduced aggregation. (A) SEC traces normalized by monomer peak height of 4NBX.B-biotin103 conjugate (biological triplicates – blue traces) and 4NBX.B WT (orange) (top panel), v119 (second panel), v149 (third panel), v186 (fourth panel) and v186_Fr (bottom panel). * indicates instrument artifact (see Figure S3C for details). (B) SPR measurement of v186_Fr/ mSA_WT_ binding (top panel) and v186_Fr/mSA_S27A_ binding (bottom panel). (C) Structural models showing sequence changes between v186 (green) and v186_Fr (cyan). Top panels show F37Y mutation in v186_Fr and potential hydrogen bonds and bottom panels show A12V mutation.

### Sequence design reduces aggregation while preserving binding affinity for the designed conjugates

4NBX.B-biotin 103 WT and v119 are well behaved by SEC with single peaks eluting at roughly the same time as the wild type 4NBX nanobody (Fig. 3A). The apparent reduced stability of v149 is consistent with its predicted higher energy score compared to v119 (Table S1). To improve the stability of v149, we explored further CDR optimizations that would better accommodate the biotin103 side chain in order to stabilize the loop and overall structure. We performed two additional rounds of CDR residue mutability analysis followed by in-parallel combinatorial designs on v149 until no further CDR mutations were predicted to be energetically favorable. Mutations accumulated in previous rounds of design were kept in subsequent rounds. In the top 20 sequences ranked by the energy score of both additional rounds of design, no additional hydrogen bonds were predicted to form with mSA, so the design with the best energy improvement, v186, was selected. The v186 variant has 6 additional CDR mutations on top of those in v149: Y101L, R107F, R56T, Y106K, D108A, and Y110S. The hydrogen bonds predicted in the v149 variant are preserved in v186 and v186 appears to bind to mSA_WT_ with a similar *K*_D_ as v149 (Fig. S5). However, SEC analysis of v186 shows even more aggregation than v149 (Fig. 3A).

Molecular dynamics (MD) simulations have been successfully applied to reveal the source of unexpected or undesired properties in designed proteins [37]. In order to explore the flaws of the current designs and inform a design strategy going forward, we perform MD simulation of 4NBX.B-biotin103 v186 in complex with mSA_WT_. The simulations showed that the CDR3 loop separates from the β-barrel framework region (supplementary movie) suggesting that optimizing the interactions of CDR3 with the framework region could lead to improved designs. We performed framework sequence design on v186. The top-ranked variant, v186_Fr, was predicted to form additional hydrogen bonds with CDR3 residues through the F37Y mutation (Fig. 3C, Top). In addition, the A12V mutation shows increased hydrophobic shielding of the β-barrel core (Fig. 3C, Bottom). Interestingly, when the same framework sequence design was performed on v149, the parent of v186, the A12V and F37Y mutations were predicted to destabilize v149, suggesting that the v186 mutations are required to realize the benefit of the A12V and F37Y mutations (Table S1-2).

4NBX.B-biotin103 v186_Fr showed significantly reduced aggregation by SEC. Collected main peak fractions did not show signs of re-aggregation when reanalyzed by SEC (Fig. 3A, S3B). SPR measurements showed the *K*_D_ of v186_Fr to be 0.20±0.03 nM, preserving the >20-fold *K*_D_ improvement observed with v149 over biotin/mSA_WT_ (Fig. 3B top panel, 5A). The kinetics profile of v186_Fr binding to mSA_WT_ was also similar to v149 (Fig. 3B top panel, 5A). When binding to mSA_S27A_, v186_Fr showed a *K*_D_ of 54±3 nM, indicating an ∼20-fold improvement resulting from both improved association and dissociate rates (Fig. 3B bottom panel, 5B).

To further investigate the structural features of v186 and v186_Fr, we performed additional 100 ns MD simulations of v186 and v186_Fr bound to mSA_WT_. During the simulations both the overall geometry of the complexes and the conformation of the biotin103 side chain remained constant with backbone-atom(?) RMSDs of less than 1.5 Å (Fig. S4, 4B first panel). The 4NBX.B nanobody scaffold has two notable solvent-inaccessible clusters of hydrophobic residues one located in the β-barrel core and the other shielded from solvent by the CDR3 loop (Fig. 4A). Buried hydrophobic surface area is generally correlated with protein folding stability, while hydrophobic surface patches and the exposure of buried hydrophobic residues are correlated with loss of stability and aggregation [9,38]. The MD simulations show reduced solvent-accessible surface area exposure for the two clusters of hydrophobic residues for v186_Fr relative to v186 supporting the idea that the F37Y and A12V mutations in v186_Fr reduce aggregation by stabilizing the framework region and reducing the exposure of hydrophobic surface area (Fig. 4A).

**Figure 4.**
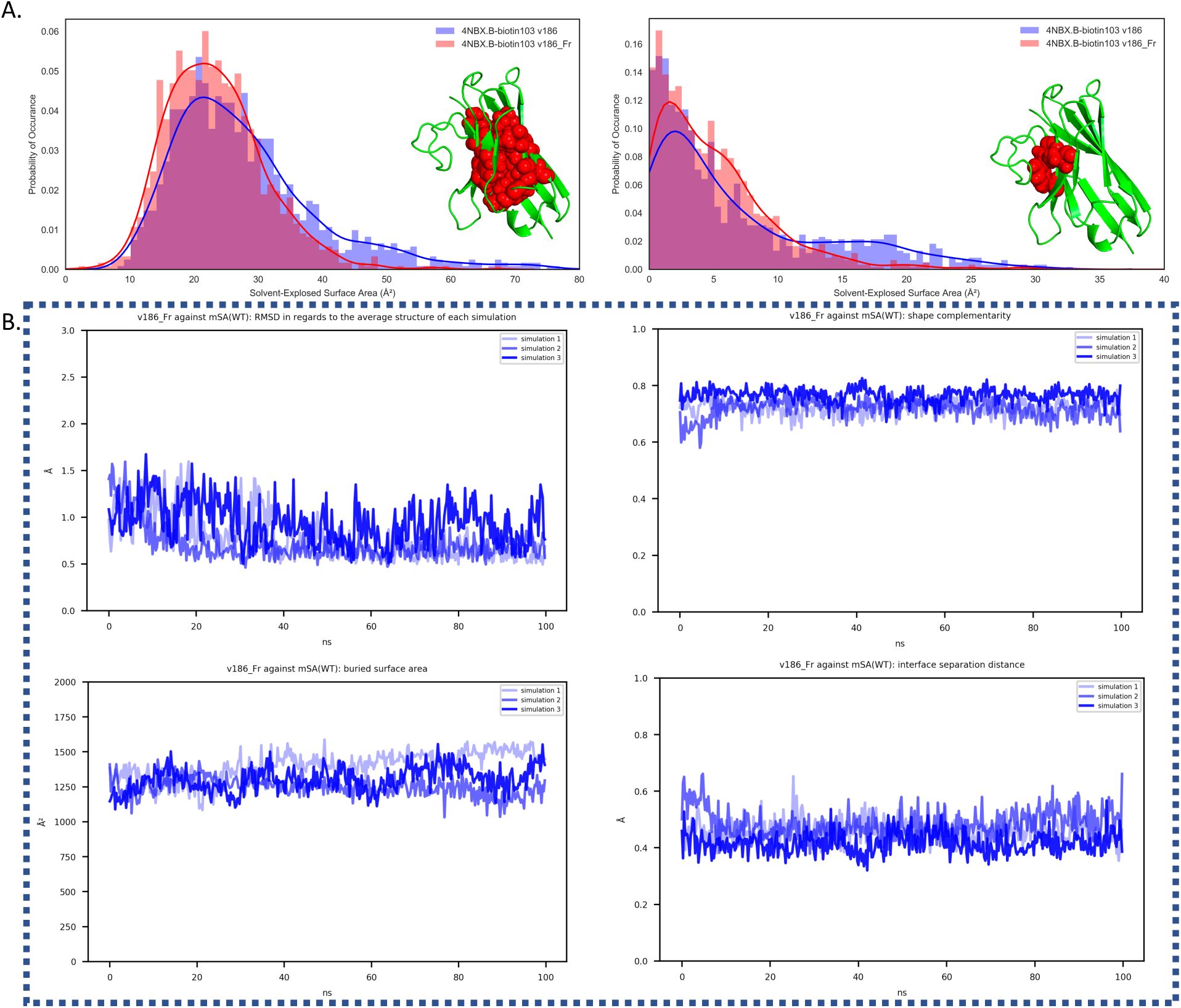
MD simulation analysis of various models. (A) Solvent accessible surface area distribution from 100 ns MD simulations for two hydrophobic residue clusters shown as red spheres in the left and right panels. Data for v186 shown in blue; data for v186_Fr shown in red. (B) Analysis of backbone RMSD (upper left panel), interface shape complementarity (upper right panel), interface buried surface area (lower left panel) and interface separation distance (lower right panel) from 100 ns MD simulations shown in triplicate for the v186_Fr/mSA_WT_ complex.

To ensure that the prepared conjugates contain a single biotin-maleimide conjugation per nanobody molecule, we used intact-protein mass spectrometry (MS) to analyze a set of variants. The most abundant species from deconvolution of the mass spectra had molecular weights within 3 Da of the expected values of mono-biotin conjugation (Fig. S6). Subpopulations within ± 20 Da of the expected molecular weights were also observed, which could result from succinimide ring opening or ion adducts (Fig. S6).

### Designing directly against mSA yields tight binding nanobody small-molecule conjugates

Because the 4NBX.B nanobody scaffold described above was not obtained by directly docking nanobody scaffolds against mSA, we performed the docking, filtering, and rotamer screening steps using mSA and selected scaffold 2X89.A with biotin conjugated to position 57 (Fig. S7A and B top panel). Similar to the 4NBX.B-biotin103 variants, the selected pose of 2X89.A was predicted to interact with mSA through a R-E interaction (in addition to other potential intermolecular H-bonds) (Fig. S7B top panel). In order to ensure a single conjugation site, two cysteines at positions 33 and 104 that form an intra-CDR disulfide bond in 2X89.A were replaced by alanine residues. The initial conjugate, 2X89.A-CCAA-biotin57 binds to mSA_WT_ with a *K*_D_ of 0.8±0.2 nM (Fig. 5A and S7B bottom panel). 2X89.A-CCAA-biotin57 was shown to aggregate by SEC analysis (Fig. S7C). To reduce aggregation, we constructed a sequence design pipeline that alternates between CDR and framework sequence optimization based on our previous experience designing 4NBX.B-biotin103 variants, and applied the pipeline to 2X89.A-CCAA-biotin57 (Fig. S7D). Six rounds of CDR sequence design and one round of framework sequence design resulted first in worsening and then ultimately improving the aggregation profile after 18 mutations were accumulated (Fig. S7E), a trend similar to what we observed in the optimization of 4NBX.B-biotin103. The resulting *K*_D_ of 2.8 nM is reduced relative to the initial design, suggesting a possible tradeoff between improved protein stability/aggregation and binding affinity.

**Figure 5:**
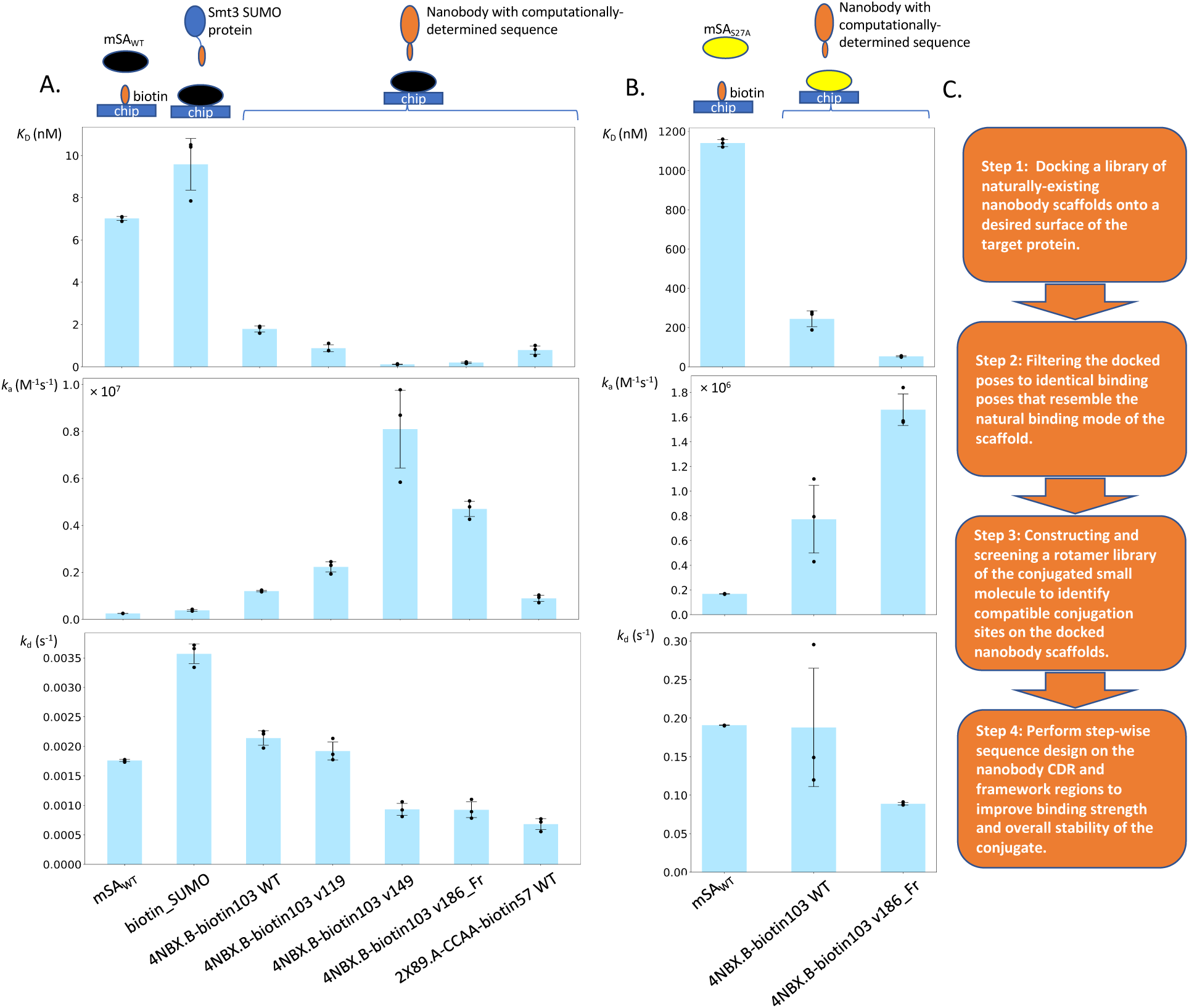
Summary of results and design workflow. (A and B) Summary of SPR determined binding affinities and kinetic parameters for the various complexes studied in this report. Setups for the SPR binding experiments are depicted by the cartoon above the lanes Individual data points are from biological triplicates. Error bars in figures show standard deviation. (C) Summary of design workflow for creating synergistically- binding nanobody-small molecule conjugates.

## Conclusion

We have demonstrated for the first time a computationally driven bridge between the worlds of small molecules and biologics by designing CDRxAbs based solely on structural information of the small molecule binding to its target. Using mSA/biotin as a model system, we showed that biotin/nanobody conjugates bind tighter to the mSA target than biotin alone. Our results showed that the affinity, kinetics and stability of the conjugates can be computationally designed and improved in a step-wise manner, which allows for a highly tunable optimization process. This approach allows the target-binding properties of the small molecule to be tuned through changes in the amino acid sequence of the conjugated antibody. The binding interface for the designed conjugates leverages the deep binding pockets common for small molecules and the broad contact interface common for antibodies [39,40]. Combining these binding characteristics into a single molecular entity may offer solutions to challenges such as off-target binding of small molecule drugs and tackling undruggable targets in pharmaceutical development.

Testing the design strategy introduced in this study on therapeutically-relevant targets is an obvious next step. In addition, examining the use of virtually-docked small-molecule/target complexes and explicitly testing for improvement of binding specificity promise to be interesting avenues for future research. There are a number of computational approaches that could be used to improve the overall design strategy. Virtually recombining structural fragments was reported to improve affinity maturation of computationally designed antibodies [26,41,42]. Algorithms that explicitly bias for the formation of hydrogen-bonding networks were also proven to be useful for improving the binding affinity and specificity of designed protein/protein interfaces [43]. Advanced loop-modeling methods and ensemble design could also facilitate a more accurate assessment of binding poses for the conjugates and potentially improve the engineering of binding specificity [44,45].

## Materials and Methods

### Computational design workflow for nanobody-biotin conjugates

A detailed description of the computational design workflows for the nanobody-biotin conjugates is included in the supplementary material.

### Plasmids, expression cell lines, and cloning of protein variants

pRSET-mSA was a gift from Sheldon Park (Addgene plasmid # 39860) [29]. The mSA S27A mutation was created by site-directed mutagenesis using commercially-available kits (NEB). 4NBX.B_C103 and 2X89.A_CCAA_C57 sequences were directly ordered from IDT, and cloned into a pHen6c vector by Gibson assembly using commercially-available reagents (NEB) [46]. The assembled pHen6c vectors harbor a PelB N-terminal signal sequence, allowing for bacterial periplasmic expression [47]. Variants of 4NBX.B_C103 and 2X89.A_CCAA_C57 were created by mutagenic PCR and assembled into a pHen6c vector by Gibson assembly using commercially-available reagents (NEB) [46]. 4NBX.B WT sequence with C103A mutation was created by site-directed mutagenesis using commercially available kits (NEB). Smt3 SUMO protein with an N-terminal cysteine was created from wild type Smt3 SUMO by mutagenic PCR, and subcloned into a pY71A(lc) vector by Gibson assembly using commercially-available reagents (NEB) [46].

### Expression and purification of wild-type mSA streptavidin and its S27A variant

Expression, purification, and refolding of mSA variants followed published protocols with slight variations [29]. The expression plasmids were first transformed into *E. coli* BL21-Gold (DE3) chemically competent cells (Agilent), which were then grown overnight in LB with 100 µg/mL of ampicillin (amp100) at 37 °C on a shaker running at 250 rpm. 1 mL of the overnight culture was used to inoculate 300 mL of TB medium (2.3 g KH_2_PO_4_, 16.4 g K_2_HPO_4_, 12 g tryptone, 24 g yeast extract, 4 mL glycerol, dissolved in water to 1 L) supplemented with 2 mM MgCl_2_, 0.1% glucose, and amp100. Inoculation was done at 37 °C and 250 rpm until an OD_600_ of 1.5 to 2. Expression was induced by 1 mM IPTG at 28 °C and 250 rpm for 18 hours. Cells were then centrifuged at 4500 g for 15 minutes at 4 °C and protein extraction was then performed using 50 mL of chemical lysis buffer. The lysis buffer was composed of 1x PBS (137 mM NaCl, 2.7 mM KCl, 10 mM Na2HPO4, 1.8 mM KH2PO4, pH 7.4), 1x CelLytic B reagent (Sigma), 0.02 mg/mL DNase1, 0.2 mg/mL lysozyme, and 1 mM protease inhibitor AEBSF (Sigma). Pellets were resuspended in lysis buffer and nutated for 4 hours at room temperature. Cell lysate was then centrifuged at 15000 g for 30 minutes at 4 °C, and the pellets was used to refold and purify the protein.

Pellets were first resuspended into 3 mL of 6 M guanidine hydrochloride in 1x TBS (50 mM Tris, 150 mM NaCl, pH 8.0) and incubated under 37 °C for 30 minutes to solubilize the proteins. Un-dissolved materials were cleared by 15000 g centrifugation for 5 minutes at 4 °C. Supernatants were then chilled on ice before dropwise addition into 40 mL of pre-chilled refolding buffer (50 mM Tris-HCl, 150 mM NaCl, 0.3 mg/mL D-biotin, 0.2 mg/mL oxidized glutathione, 1 mg/mL reduced glutathione, pH 8.0) while stirring. The refolded mSA protein was then allowed to incubate on ice for an additional 2 hours with stirring before centrifugation at 15000 g for 30 minutes at 4 °C to remove insoluble material. The supernatants were then supplemented with 20 mM imidazole and then loaded onto 1 mL bed volume of Ni-NTA agarose beads (Qiagen) that were pre-washed with 5 column volumes of 1x PBS. Sample loading was performed by gravity flow. The column was then washed with 10 column volumes of 1x PBS supplemented with 20 mM imidazole before 3 mL of 1x PBS supplemented with 500 mM imidazole was used to elute the protein. The entire Ni-NTA purification process was performed at 4 °C. The 3 mL of purified protein was then dialyzed against fresh 1 L of 1x PBS (pH 8.0) at 4 °C 3 times using Slide-A-Lyzer 10 kDa molecular-weight cutoff dialysis cassettes (Thermo). Each dialysis step was performed for at least 4 hours. The dialyzed mSA product was further purified at 4 °C using a Superdex 75 10/300 GL SEC column (GE) with 1x PBS (pH 7.4) as running buffer. The fractions corresponding to monomeric mSA were collected for subsequent experiments.

### Expression and purification of Smt3 SUMO protein with N-terminal cysteine

The expression plasmid was first transformed into *E. coli* BL21-Gold (DE3) chemically competent cells (Agilent), which were then grown overnight in LB with amp100 and 1% glucose at 37 °C and 250 rpm. 1 mL of the overnight culture was then used to inoculate 300 mL of TB medium with 2 mM MgCl_2_, 0.1% glucose, and amp100 at 37 °C and 250 rpm until an OD_600_ of 1.5 to 2. Expression was then induced by 1 mM IPTG at 28 °C and 250 rpm for 18 hours. Cells were then pelleted at 4500 g for 15 minutes at 4 °C, resuspended in 50 mL of chemical lysis buffer supplemented with 5 mM 2-mercaptoethanol (BME) and incubated for 4 hours at room temperature to release the expressed protein. Lysate supplemented with 20 mM imidazole was cleared by centrifugation at 15000 g at 4 °C for 30 minutes and the supernatant was then loaded onto 1 mL bed volume of Ni-NTA agarose beads (Qiagen) pre-washed with 5 column volumes of 1x TBS (pH 7.3). Sample loading was performed by gravity flow. The loaded column was then washed with 5 column volumes of 1x TBS (pH 7.3) supplemented with 20 mM imidazole and 5 mM BME and another 5 column volumes of 1x TBS (pH 7.3) supplemented with 20 mM imidazole. 3 mL of 1x TBS (pH 7.3) supplemented with 500 mM of imidazole was used to elute the proteins. All Ni-NTA purification procedures were performed under 4 °C. Purified protein was then concentrated to ∼0.5 mL using Amicon 10 kDa molecular-weight cutoff centrifuge filters (GE), and stored for subsequent experiments.

### Expression and purification of nanobodies

The pHen6c expression plasmids were first transformed into *E. Coli* BL21-Gold (DE3) chemically competent cells (Agilent), which were then grown overnight in LB with amp100 and 1% glucose at 37 °C and 250 rpm. 1 mL of the overnight culture was then used to inoculate 300 mL of TB medium with 2 mM MgCl_2_, 0.1% glucose, and amp100 at 37 °C and 250 rpm until an OD_600_ of 1.5 to 2. Expression was then induced by 1 mM IPTG at 28 °C and 250 rpm for 18 hours. Cells were then pelleted under at 4500 g for 15 minutes at 4 °C and resuspended in 12 mL of TES periplasmic extraction buffer (0.2 M Tris, 0.5 mM EDTA, 0.5 M sucrose, pH 8.0) (supplemented with 5 mM BME if the nanobody had a cysteine handle for conjugation) before incubation on ice with shaking at 32 rpm for 1 hour [47,48]. 18 mL of 4x diluted TES buffer (supplemented with 5mM BME if the nanobody had a cysteine handle) was then added to the cells which were incubated on ice at 32 rpm for another hour [48,49]. After periplasmic extraction, the cells were pelleted at 15000 g at 4 °C for 30 minutes and the supernatant was supplemented with 20 mM of imidazole before loading onto 1 mL bed volume of Ni-NTA agarose beads (Qiagen) pre-washed with 5 column volumes of 1x TBS (pH 7.3). Sample loading was performed by gravity flow. The loaded column was then washed with 10 column volumes of 1x TBS (pH 7.3) supplemented with 20 mM of imidazole. 3 mL of 1x TBS (pH 7.3) supplemented with 500 mM of imidazole was used to elute the proteins. For nanobodies with a cysteine handle, 5 mM BME was added to the first 5 column volumes of wash buffer and the elution buffer was supplemented with 5 mM TCEP. The elution buffer was incubated with Ni-NTA beads for 30 minutes before eluting the proteins. The entire Ni-NTA purification process was performed at 4°C. Purified nanobodies with a cysteine handle in 3 mL of elution buffer were concentrated to ∼0.5 mL using Amicon 10 kDa molecular-weight cutoff centrifuge filters (GE) and stored for subsequent experiments. 4NBX.B WT was instead further purified using a Superdex 75 10/300 GL SEC column (GE) with 1x PBS (pH 7.4) as a running buffer and the fractions corresponding to the monomeric protein were collected for subsequent experiments.

### Biotin C2 maleimide conjugation and purification of conjugates

Maleimide labeling on the solvent exposed cysteines of nanobodies followed a published protocol with some modifications [50]. Purified nanobodies from storage were first incubated with another 5 mM TCEP supplement at 4 °C for 2 hours and then buffer exchanged into 1x TBS (pH 7.3) using HiTrap desalting columns (GE) at room temperature to remove TCEP. Thawed stock solutions (100 mM in DMSO) of biotin C2 maleimide (AnaSpec) were immediately added to the buffer-exchanged nanobodies to final concentration of 1 mM before the reaction mixture was nutated at 4 °C for 4 hours with a tinfoil cover to avoid light contact. Excess maleimide stock solutions were discarded. Biotin C2 maleimide was in >20-fold molar excess over the nanobody in the reaction mixture. The reaction mixture was then filtered using 0.2 µm syringe filters (Thermo) to remove precipitated protein and then buffer exchanged to 1x TBS (pH 7.3) using PD-10 desalting columns (GE) at room temperature to remove excess maleimide reagent. The labeled nanobodies were further purified at 4 °C using a Superdex 75 10/300 GL SEC column (GE) with 1x TBS (pH 7.3) as the running buffer and the fractions corresponding to monomeric protein were collected for subsequent experiments. Maleimide labeling of Smt3 SUMO protein with an N-terminal cysteine followed the identical procedure described above.

### Intact protein mass spectrometry (MS) workflow to analyze conjugation efficiency

HPLC-MSD (HP, Agilent) was used to assess the labeling efficiency of the nanobody-biotin conjugates. Conjugates were first dried using a spin vacuum evaporator and then resuspended in 0.2% formic acid. A C3 HPLC column was used to purify the protein sample before MS analysis. Before running samples, the column was first washed with isopropyl alcohol (IPA). Analysis of the conjugation efficiency of 4NBX.B-based conjugates was performed by deconvoluting each eluted sample HPLC peak using the following parameters: positive adduct ion +H 1.0079 Da, negative adduct ion –H -1.0079 Da, molecular weight cutoff 5000-80000 Da, maximum charge 90, minimum peaks 5, ion PWHH 0.6 Da, molecular weight agreement 0.05%, noise cutoff 0, abundance cutoff 10%, molecular weight assignment cutoff 40%, and envelope cutoff 50%. Deconvolution was performed with ChemStation (Agilent). For each sample of interest, approximately 0.1 to 1 µg of material was used for the analysis.

### Surface plasmon resonance (SPR) analysis of binding affinity and kinetics

A Biacore T200 instrument (GE) was used to perform SPR analysis. 4NBX.B WT, 4NBX.B-biotin103 conjugates, and 2X89.A-CCAA-biotin57 conjugates were first buffer-exchanged into HBS-EP+ buffer (Teknova) using Amicon 10 kDa molecular-weight cutoff centrifuge filters (GE). Protein concentration was then determined by BCA assay using commercially-available kits (Thermo). The calibration curve for the BCA assay was prepared using purified 4NBX.B WT, which was also buffer exchanged into HBS-EP+ but had concentrations determined by A_280_ readings using an extinction coefficient 30035 M^-1^cm^-1^. For SPR analysis, biotin pentylamine (Thermo), mSA_WT_, and mSA_S27A_ were separately immobilized on CM5 censor chips (GE) using an EDC/NHS amine coupling kit following standard protocols (GE). Binding kinetics were measured by single-cycle kinetics experiments. Biotin pentylamine was immobilized at 7.5 mM concentration to reach a target surface density of ∼200 resonance units (RUs) [51]. In order to compare how binding events changed in response to different surface densities, surfaces with three different densities of immobilized mSA_WT_ at ∼1000 RU, ∼2500 RU, and ∼3000 RU were prepared under immobilization concentrations 0.1 µM, 0.5 µM, and 1 µM, respectively. Immobilization of mSA_S27A_ was also performed at 0.01 µM and 0.05 µM concentrations with target surface density of ∼200 and ∼600 RU. The fitted affinities and kinetics of identical conjugates using different surfaces densities were not significantly different. Reference channels were either treated with EDC/NHS using HBS-EP+ buffer alone or with 1 µM 4NBX.B WT to assess non-specific interactions. No non-specific interactions were observed. All immobilization samples were dissolved in acetate buffer (pH 4.5).

Binding experiments were performed at 25 °C. HBS-EP+ was used as running buffer. The flow channels were first incubated in the running buffer before analytes at 5 different concentrates were consecutively injected at a flow rate of 30 µL/min through both the reference channel and the sample channel containing the immobilized molecules of interest. After injections, the bound analytes were allowed to dissociate for 10 minutes in order to generate dissociation curves. HBS-EP+ buffer was then washed through both the reference and sample channels in order to regenerate the surfaces for the next binding experiments. Sensorgrams were processed with Biacore evaluation software using a 1:1 kinetics model. For immobilized mSA_WT_, global fitting of bulk shifts was turned on to accommodate the observed bulk shifts before and after each injection event.

### Molecular dynamics (MD) simulation protocols

Molecular Dynamics simulations were carried out using ACEMD (Acellera) [52]. Each system was placed in a box with dimensions selected to allow 12 Å of clearance on each side. The systems were solvated using the TIP3P water model [53] and counter ions were added to bring the net charge to zero. The systems were then minimized for 500 steps and equilibrated for 5 ns before 100 ns production runs were carried out at 300 K. All experiments utilized the Amber ff14SB force field and a 4 femtosecond timestep [54]. Data from the equilibration run was not included in subsequent analysis, and where replicates were collected no part of the intermediate data was reused. Parameters for the biotin-CH2-CH2-succinimide-S-CH3 “side chain” were prepared using Antechamber and utilized RESP charges calculated with Gaussian 09 [55,56]. Calculation of solvent accessible surface areas was performed using MDTraj, and hydrogen bonding was assessed using a tcl script written for VMD [57, 58].

## Acknowledgements

We thank Monica C. Breckow for technical assistance; Dr. Jost G. Vielmetter and the Caltech Protein Expression Center for consultation with Biacore experimental design and data interpretation; and, Dr. Barry D. Olafson and Paul Chang from Protabit LLC for consultation on the implementation and use of the TRIAD and BIOGRAF software programs.

## Notes

### Competing Interest Statement

The authors have declared no competing interest.

