## Supplemental Information for "CDRxAbs: Antibody Small-Molecule Conjugates with Computationally Designed Target-Binding Synergy"

[c]Present address: Jingzhou Wang, Merck Research Laboratories, 213 E Grand Ave, South San Francisco, CA 94080.

This file includes:

Supplementary text
Figs. S1 to S7
Tables S1 to S4
References for SI

**Supplementary Information Text:**

**Computational design workflow for nanobody-biotin conjugates**

The protein design software package TRIAD [1] together with PyMOL (Schrodinger) and OpenBabel [2] were used to perform computational protein design and analysis in this study. The detailed process and setup are described below.

**A. Design process of 4NBX.B-derived nanobody-biotin conjugates**

Searching optimal streptavidin-binding nanobody CDR conformations by docking and loop modeling

A monomeric streptavidin model, S45A/T90A/D180A, was prepared from the crystal structure of wild-type tetrameric streptavidin (PDB ID: 1MK5). A single subunit was extracted and standardized using TRIAD [1]. The S45A/T90A/D128A mutations were then introduced using the TRIAD sequence-design module. Initially, we attempted optimizing nanobody/streptavidin binding conformations by protein-protein docking followed by loop modeling of the nanobody CDRs. We used a published nanobody structure (PDB ID: 5VNW, chain C) as the starting nanobody scaffold with all CDR residues replaced by alanine using TRIAD sequence design in a hope to avoid sequence bias [3]. The nanobody scaffold was docked onto a set of manually selected surface residues surrounding the binding pocket of the monomeric streptavidin model. Docking was performed using an FFT-based docking algorithm and the top 15 CDR binding poses were kept [4]. CDR loop modeling of each pose was then performed to attempt optimizing binding to the target using the TRIAD loop modeling module. Only the top solution for each pose was kept. Based on visual analysis, no solutions had reasonable CDR conformations that would be beneficial for binding. The sequence of the monomeric streptavidin used for these calculations is listed below with the residues selected as docking targets surrounded by brackets:

EAGITGTWY(NQLGS)TFIVTAGADGALTGT(YEAAVGNAESRY)VLTGRYDSAPATDGS GTALGWTVA(WKNNYRNAHSA)ATWSGQYVGGAEARINTQ(WLLTSGTTEANAWKSTLVGHA TFT)KVK.

<u>Searching optimal streptavidin-binding nanobody CDR conformations by docking native CDR sequences and conformations</u>

154 nanobodies crystal structures with resolutions better than 3Å, complete electron density, and structurally diverse target-binding CDR conformations were assembled from published nanobody/target complexes from the PDB (see section C below). The TRIAD surface-complementarity module was used to perform interface analysis on the 154 nanobody-target complex structures to collect statistics on separation distance, shape complementary, and buried interface area for use in evaluating designs (Fig. S1B) [5]. CDRs for each nanobody were annotated following established nanobody CDR-mapping criteria [6] but with softer edge cutoffs so that the sampled geometries of nanobody approaching are not too stringent.

Each nanobody was docked to the previously-selected binding pocket surface (defined above) using the monomeric streptavidin model and the annotated CDR loops. The top 15 poses for each docking trial were kept. Initially, we performed alanine-replacement on all CDR residues before docking. However, several CDR sequence design trials using the dock complexes resulted in the selection of mostly small amino acids, suggesting that the poly-alanine CDRs resulted in streptavidin/nanobody distances that were too close. Therefore, we decided to dock the 154 nanobody structures using their native CDR sequences. 2310 docked poses were generated and then filtered using three selection steps to identify the most native-like binding poses. Step one selected poses with interface separation distance, shape complementary score, and buried interface area within 1 standard deviation of the values obtained for the 154 nanobody/target structure dataset (Fig. S1B). This filter resulted in 231 poses. Step two then selected nanobody poses where the nanobody/streptavidin interface utilized >80% of the nanobody residues that participate in the native nanobody/target interface, where the interface residues were determined using a PyMOL script that evaluates changes in solvent accessibility between the docked and undocked structures [7]. This filter resulted in 31 poses. The last step selected poses that directly blocked the biotin binding pocket and resulted in 7 final poses (Fig. S1C). Binding pocket blockage was assessed using PyMOL to calculate the change in solvent accessible area between the bound and unbound complex for the following selected residues that are a subset of the docking target residues and closely surround the biotin molecule (highlighted in parentheses along with the full streptavidin sequence):

EAGITGTWYN(QLGS)TFIVTAGADGALTGTYEAA(VGNA)ESR(Y)VLTGRYDSAPATDGS GTALGWTVA(W)KNNYRN(AHS)AATWSGQYVGGAEARINTQ(W)L(L)T(S)GTTEANAWKSTLV(G)H(A)T(FT)KVK.

<u>Rotamer library generation for biotin C2 maleimide side chain and conjugation plan determination</u>

A rotamer library for the biotin-CH2-CH2-succinimide-S-CH3 "side chain" was constructed using OpenBabel [2] and virtually screened against CDR amino acid locations on the 7 poses from above to find the optimal conjugation sites. In the rotamer library, the biotin portion remains fixed, while the CH2-CH2-succinimide-S-CH3 portion is structurally diversified through torsion angle modifications. Screening was done by measuring the distance between the terminal carbon of the rotamer and the Cβ of respective CDR residues, steric clash between the rotamer and streptavidin, steric clash between the rotamer and the nanobody, and the angle of the rotamer's terminal carbon with respective to the target Cα attachment point on the nanobody. Rotamers that were <1 Å from the target Cβ, clashed with streptavidin by <1 unit, clashed with the nanobody by <15 units, and had attachment point angles between 100-120 degrees were kept for further evaluation. Measurements were performed only for CDR residues that are originally alanine, which is similar in size to cysteine, as we hypothesized that making an alanine to cysteine mutation would be less likely to cause serious structural consequences for the nanobody. Alanine 103 on 4NBX.B was the only conjugation site that passed all of the filters, and the rotamer that clashed least with both streptavidin and the nanobody was selected for further processing. To prepare the final conjugation structure, excess atoms were removed and a bond was made between the Cβ of the biotin-CH2-CH2-succinimide-S-CH3 "side chain" and the Cα of 4NBX.B at position 103. The conjugated 4NBX.B-biotin103 structure bound to

S45A/T90A/D128A streptavidin was then relaxed using BIOGRAF [11] with force restraints placed to maintain the biotin-streptavidin hydrogen bonds and torsion angles of the aliphatic arm portion of biotin. To prepare the 4NBX.B-biotin103/mSA structure, the crystal structure of mSA (PDB ID: 4JNJ) was aligned to the modeled triple-mutation streptavidin structure, and the resulting complex with 4NBX.B-biotin103 was relaxed using BIOGRAF with the same force restraints as above [11].

4NBX.B-biotin103 sequence designs

CDR sequence design was first performed on the 4NBX.B-biotin103/mSA model. CDR residues were determined following established nanobody CDR-mapping criteria [6], and specifically by first aligning the 4NBX.B model with an example nanobody structure in agreement with that criteria (PDB ID: 5VNW, chain C), and then selecting the corresponding CDR residues on 4NBX.B. Residues 27-34 were selected as CDR1, 47-60 were selected as CDR2, and 98-111 were selected as CDR3. Single residue site-saturation mutation scans were performed on each CDR position using reduced sets of amino acids that were reported to be frequently used in each corresponding CDR position [6]. The wild-type amino acid was considered for each position. The amino acid sets for each position were as follows:

| Positions | Amino Acids |
|---|---|
| 27 | N, S, T, Y |
| 28, 51 | I |
| 29 | S, F |
| 34 | M |
| 47 | F, L |
| 48 | V |
| 49, 60, 98 | A |
| 50, 55 | A,G , S, T |
| 52 | A, D, G, Q,S , T |

| | |
|---|---|
| 54 | G |
| 56 | I, N, S,T |
| 57, 58 | T |
| 59 | N,Y |
| 60, 111 | Y |
| 109 | F, H, L,Y |
| 30-33, 53, 99-102, 104-108, 110 | A, R, N, D, Q, E, G, H, I, L, K, F, P, S, T, W, Y, V |

Position 103 with the attached biotin "side chain" was left un-designed and the coordinates for all atoms were left unchanged. The Rosetta force field [8] option with covalent terms was used in TRIAD for design the calculations. The BIOGRAF-relaxed 4NBX.B-biotin103/mSA structure was used as the input design model. During the design calculations, residues that were within 10 Å from the position being evaluated were allowed to repack. Each rotamer optimization for the position being evaluated was initiated by random rotamer configurations and then repacked while the Cα backbone was allowed to relax through Cartesian minimization to optimize the structure for different amino acid choices, which were then ranked given the energy scores of the corresponding modeled structures after iterative rotamer repacking and backbone relaxation. The chemical attributes of the biotin103 "side chain" were generated by TRIAD and then used during the design calculations of the energy scores. 10 runs with different random seeds were performed for each design calculation, and averaged to reflect the final amino acids preference for each site that underwent single-site mutation calculations. Mutation choices with lower Rosetta energy unit than the WT amino acids were kept as designable mutations. Designability of each site was reflected by the sum of Rosetta energy unit differences of designable mutations from the corresponding WT amino acid choice. As the result, the output designable sites ranked by designability were as follows: 105, 109, 107, 104, 106, 32, 108, 31, and 56. Those sites were also separately grouped into two bins. Bin 1 contains sites that interact with the original target of 4NBX.B: sites 105, 104, 32, and 31, of which the order was ranked by designability. Bin 2 contains sites that do not interact with the original target of 4NBX.B: sites 109, 107, 106, 108, and 56, of which the order was ranked by designability.

Combinatorial designs with different choices of designable sites were performed in parallel. Combinatorial design 1 was performed on all the 9 designable sites. Combinatorial design 2 was performed on the designable sites in bin 1 only. Combinatorial design 3 was performed on the designable sites in bin 1 and bin 2, with an exception that for bin 2 sites, only mutation choices that are different from WT amino acid with >1 Rosetta energy units were used. Combinatorial design 4 was performed on the designable sites in bin 1 and bin 2, with an exception that for bin 2 sites, only the top-ranked mutation choice by energy score was used.

Combinatorial designs were performed with the same configurations as single-site mutation designs that were described before, with one difference: the output sequences from the 10 parallel design runs were re-ranked by threading the sampled sequences in each run individually onto the backbone of the input structure, followed by rotamer repacking and backbone Cartesian minimization. The TRIAD-modeled structures and Rosetta energy scores of the top 20 sequences of the re-ranked sequences were used to evaluate design results.

Structures of the top 20 sequences for the 4 combinatorial designs were analyzed by PyMOL to identify intermolecular H-bonds between $mSA_{WT}$ and 4NBX.B-biotin103 variants, and intramolecular H-bonds within 4NBX.B-biotin103 variants, using a publically-available PyMOL script that relies on the "find_pairs" command module of PyMOL [9]. The goal was to find sequences with improved overall energy score, new intermolecular H-bonds with $mSA_{WT}$, and intramolecular H-bond profile comparable to 4NBX.B-biotin103 WT, as the imbalance of forming new interactions with targets and keeping the structural integrity was a common reason behind the failure of designing protein-protein interactions [10]. All combinatorial designs output sequences with improved energy scores, but only combinatorial design 2 output the top 20 sequences with an average number of intermolecular H-bond higher than that of the 4NBX.B-biotin103 WT against $mSA_{WT}$. The top 20 sequences in combinatorial design 2 also had the highest average number of intramolecular H-bonds in the nanobodies among the 4 designs. Variant v119 was the top-ranked sequence in combinatorial design 2, and variant v149 had the highest number of predicted intermolecular H-bonds among the top 20 sequences (Table S1).

To improve the stability of v149, we hypothesized that a suitable method would optimize the protein structure while keeping the designed interactions contributed by R31/S32/A104/R105, the new anchoring spots, unchanged, without altering the target backbone structure too much. Therefore, we devised a sequential design workflow that creates stepwise local structural optimizations that compensate for the mutations built-up in previous steps. As the result, subsequent rounds of CDR sequence designs were performed on v149. For each round, single-point mutation scan was first performed on the CDR residues using the identical setup as the first round of design. The calculation results were processed in the same way as the first round of design, with an exception that only mutation choices that are different from WT amino acid with >1 Rosetta energy units were kept for all further combinatorial design calculations. Next, skipping the sites that were mutated in previous rounds of design, four combinatorial designs were performed on the designable sites reported by the single-point mutation scan calculation. Combinatorial design 1 was performed on all designable sites with the reported designable mutation choices. Combinatorial design 2 was performed only the designable sites in bin 1 with the reported designable mutation choices. Combinatorial design 3 was performed on top 5 designable sites with the reported designable mutation choices. Combinatorial design 4 was performed on 5 designable sites ranked by designability, but with a bias on sites in bin 1. In other words, sites in bin 2 were not used unless the number of sites in bin 1 was smaller than 5. All combinatorial design calculations were performed and processed under the same setup as the first round of design.

The second round of design was performed using the output structure of v149 from the first round of design as input. No improvement in the number of intermolecular H-bond formation was observed for the outputs of all the 4 combinatorial designs, while the numbers of intramolecular H-bond were minimally different among the designs. Therefore, the design result with the biggest overall difference in energy score among the top 20 sequences against v149, combinatorial design 4, was chosen, and of which the sequence with the best energy score, v149 plus Y101L/R107F, was selected as the input structure for the third round of design. Combinatorial design 4 was performed with sites 27/59/101/107/110. In the third round, again no improvements in the number of intermolecular and intramolecular H-bonds were observed among the 4 combinatorial designs. So, the sequence with the best energy score, v186 (v149 plus Y101L/R107F/R56T/Y106K/D108A/Y110S), of combinatorial design 1 whose top 20 sequences showed biggest overall improvement in energy scores against the input sequence was selected. Combinatorial design 1 was performed with sites 27/29/56/106/108/110. A further round of CDR design was performed on v186 and all combinatorial design results returned sequences with worse energy score than v186.

Because v186 turned out to be even more prone to aggregation than v149, and based on MD simulation results of v149 against $mSA_{WT}$, we hypothesized that only mutating CDRs was not sufficient. Therefore, we proceeded to design the framework regions of v149. Because the frameworks of nanobodies are highly conserved [6], a suitable sets of amino acid choices and locations would be crucial for the design calculation. Because the previous CDR designs were based on a published summary of nanobody CDR sequence diversity, we referred to the framework sequence used in that study for framework sequence design [6]. We aligned 4NBX.B-biotin103 v186 with chain C of 5VNW, identified framework sites where the two nanobodies differ, and performed a combinatorial design with the selected sites being one or the other amino acid choice. Site positions and sequence choices were as follows: 5-V/G, 12-A/V, 35-A/G, 37-F/Y, and 40-P/A. The v186 structure output by the third round of design was used as input, and the configurations and processing of design calculation were the same as the combinatorial CDR sequence designs described previously. The top-ranked sequence by energy score was v186_Fr (v186 plus A12V/F37Y) (Table S2). Framework design with identical amino acid sites, sequence choices, and calculation configurations was performed using v149 as input, and the design results were also reported in this study for comparison (Table S3).

As a comparison, we performed two rounds of CDR sequence design using the above-described configurations on v119. Combinatorial designs in both rounds failed to produce variants with new inter-molecular H-bond formation in the top 20 variants, so the combinatorial designs with best overall energy improvement than the input sequence were chosen, and the top-ranked sequences by energy score in those designs were used as input for further rounds of design and experimental testing. No improvements in kinetics and affinity were observed in these selected sequences (data not shown), in agreement with the unchanged inter-molecular H-bond profiles and with what we observed for v186 versus v149.

**B. Design process of 2X89.A-derived nanobody-biotin conjugates**

Docking, pose filtering, rotamer screening, and binding pose generation for 2X89.A-CCAA-biotin57 WT against $mSA_{WT}$

The 154 nanobody structures from PDB with native CDR sequences were docked against a manually-selected set of surface residues around the biotin-binding pocket of mSA, in the same way as described in part A. The amino acids being docked against are highlighted below by parentheses with the rest of mSA sequence shown for reference:

GAEAGITGTWYN(QSG)STFTVTAGADGNLTGQY(ENRAQGTG)C(QNSP)YTLTGRYNGTKLEWRVEWN(NSTENCH)SRTEWRGQYQGGAEARINTQWNLT(YEGGSGPATEQGQDT)FTKVK.

Filtering procedures of the docked poses were also identical to those introduced in part A, with an exception that the following residues (highlighted by parentheses) were selected as the target for binding pocket blockage filter:

GAEAGITGTWYN(QS)GSTFTVTAGADGNLTGQYENRAQGTGCQNSPYTLTGRYNGTKLEWRVEWNNSTENCHSRTEWRGQYQGGAEARINTQWNLT(YEGGSGPATEQGQDT)FTKVK.

6 poses that respectively comprised nanobodies 2X89.A, 3EBA.A, 4LHQ.B, 4OCL.C, 3V0A.C, and 4P2C.G passed the series of filters. Visual inspection of the poses revealed that the 4OCL.C and 3V0A.C binding poses showed significant contacts that are mediated by nanobody residues outside of the CDRs, so the corresponding two poses were discarded since these binding modes were potentially unrealistic. As the result, 4 final binding poses were kept for the evaluation of optimal conjugation sites (Fig. S7A).

The biotin-CH2-CH2-succinimide-S-CH3 rotamer library built in part A was used again for rotamer screening. Because the design process of the 4NBX.B-derived CDRexAbs demonstrated that our design capability allowed structural stability of the conjugates to be designed after binding synergy was designed, we did not put much emphasis on preserving structural integrity in the early stage of the design process of 2X89.A-derived CDRexAbs. Therefore, instead of screening only against alanine CDR residues, all CDR residues were screened by measuring the distance between the terminal carbon of the rotamer and the Cβ of respective CDR residues, steric clash between the rotamer and the streptavidin, steric clash between the rotamer and both proteins, and the angle of the rotamer terminal carbon approaching the respective attachment spot. Conjugation geometries that clashed with streptavidin by <0.5 unit, clashed with both proteins by <10 units, approached the attachment spot by 100-120 degree, and were <2 Å away from the Cβ of screened conjugation sites were kept. Only one rotamer that was screened against I57 of 2X89.A passed the filter. The final conjugation structure was prepared by Biograf, under the same parameters as described in part A [11].

To remove the intra-CDR disulfide bond in the Biograf-relaxed structure, C33A/C104A mutations were introduced by TRIAD sequence design module to create the finalized model of 2X89.A-CCAA-biotin57/mSA for further sequence design optimization.

Subsequent sequence design of 2X89.A-CCAA-biotin57 conjugates

Summarizing the experience from the design process of the 4NBX.B-biotin103 conjugates, we gained the following insights into the sequence design principles of CDRexAbs:

1. Performing sequential rounds of design on limited sets of amino acid sites and choices that are recommended by iterative energetic and structural analysis allows functionally-improved CDRexAb mutants to be discovered without experimentally screening a large set of sequences.

2. New intermolecular interactions between the nanobody scaffold and the target can be engineered first before further mutations are introduced to optimize the structural integrity of the conjugates.

3. Simply mutating CDR residues is not sufficient for structural optimization of the conjugates, and mutated CDR residues likely need accommodation by introducing mutations in the β-barrel framework region.

Based on the above principles, we established a rudimentary sequence design pipeline, and tested it on 2X89.A-CCAA-biotin57 to create mutants that are less prone to aggregation. The pipeline is detailed below (Fig. S7D):

1. The pipeline performs sequential rounds of sequence design that is either restricted on CDR residues or framework residues. The first round of design is performed on CDR residues. Residues that are mutated in previous rounds are kept from mutation in further rounds.

2. H-bonds are still the only intermolecular interactions that are explicitly evaluated after design calculations and biased towards for sequence selection, but other types of interactions can be evaluated if they are deemed to be crucial for specific scenarios.

3. CDR sequence design follows the procedures of designing the CDR loops of v149, as described in part A, with one exception: besides the four combinatorial designs, it is optional that two additional combinatorial designs can be performed in parallel, respectively on all identified sites and amino acid choices in bin 2, and on the top 5 designable sites with a bias on sites in bin 2. All combinatorial designs are evaluated together as described in part A.

4. Framework design follows the procedures of designing the framework of v186, as described in part A.

5. To evaluate the design results of CDR design, the following steps are used:

a). Sequences that showed worse energy score than the immediate parent sequence are discarded.

b). Sequences with the number of intermolecular H-bonds lower than the immediate parent sequence are discarded.

c). If no sequences survived filters a and b, perform framework design on the immediate parent sequence.

d). For sequences pass the filters, the sequence that has the highest number of intermolecular H-bonds and the best energy score among sequences that share the same number of intermolecular H-bonds is kept as input for the next round of CDR design.

6. To evaluate the design results of framework design, the following steps are used:

a). Sequences that showed worse energy score than the immediate parent sequence are discarded.

b). Sequences with the number of intermolecular H-bonds lower than the immediate parent sequence are discarded.

c). Sequences with the number of nanobody intramolecular H-bonds lower than the immediate parent by >1 are discarded.

d). If no sequences survived filters a-c, design fails.

e). For sequences pass the filters, the sequence that has the highest energy score is kept as input for the next round of CDR design.

The outputs after rounds 3, 5, 6, and 7, which are variants v37, v42, v20, and v5 were selected for experimental testing.

**C. Nanobody sequences that are used in this study for docking and binding pose selection**

Note: Sequences are represented by [PDB ID].[chain name]

1BZQ.K, 1JTT.A, 1KXQ.E, 1OP9.A, 1ZVH.A, 1ZVY.A, 2X89.A, 3EBA.A, 3JBC.7, 3JBE.7, 3JBF.7, 4GRW.E, 4I0C.C, 4W6W.B, 5BOP.A, 5C2U.B, 5FOJ.A, 5M13.B, 5TJW.K, 2XT1.B, 3K74.B, 3SN6.N, 4C57.C, 4EIZ.C, 4FHB.D, 4GRW.F, 4HEM.E, 4KML.B, 4LGP.B, 4N9O.B, 4NBY.B, 4NBZ.B, 4TVS.b, 5C3L.D, 5F1K.C, 5F1O.B, 5H8O.A, 5IMK.B, 5J57.B, 5JA8.B, 5JMO.C, 5KU2.7, 5LHN.B, 5NBD.C, 5O03.C, 5USF.C, 5UZ7.N, 5VXL.B, 5VXM.B, 4Z9K.B, 5IVN.A, 3QXT.A, 4DK3.A, 4GFT.B, 4KRL.B, 4LHQ.B, 4N1H.B, 4S10.A, 5OJM.K, 5UKB.a, 5VXK.B, 1I3U.A, 1RJC.A, 4GRW.H, 4IOS.D, 4OCL.C, 5BOZ.G, 5JDS.B, 5M2M.D, 1KXV.C, 1QD0, 1RI8.A, 1ZV5.A, 3JBD.7, 3K3Q.A, 3QXV.A, 3RJQ.B,

3STB.A, 4AQ1.B, 4CDG.C, 4I13.B, 4QO1.A, 4W6X.B, 4WEM.B, 4XT1.C, 5E0Q.A, 5G5R.B, 5GXB.B, 5LHR.B, 5LWF.C, 5O8F.K, 5OCL.B, 4C58.B, 2XXM.B, 3JBG.7, 4X7F.C, 5F21.B, 5F7K.C, 5HVG.B, 4LHJ.B, 4M3K.B, 5HVF.B, 5MJE.B, 1G6V.K, 2X6M.A, 3K1K.C, 3V0A.C, 4U3X.A, 5E5M.B, 5HGG.S, 5HM1.A, 5J56.B, 5JA9.A, 5MWN.N, 5O2U.B, 5OVW.G, 5TOK.D, 1KXT.B, 3CFI.C, 3K81.A, 4EIG.B, 4HEP.G, 4LGR.B, 4MQS.B, 4NBX.B, 4NC2.B, 4P2C.G, 4W6Y.B, 4YGA.B, 5DFZ.E, 5IP4.A, 5J1S.C, 5JQH.C, 5KTZ.7, 5KU0.7, 5KWL.7, 5L21.B, 5MY6.B, 5O02.C, 5O04.E, 5O0W.E, 5OCL.A, 5OMN.C, 5TOJ.D, 5UK4.a, 5VXJ.B, 4X7C.C, 3EZJ.B, 5F7L.B, 5M30.D, 5M94.B, 3P0G.B, 4LDE.B, 2BSE.D

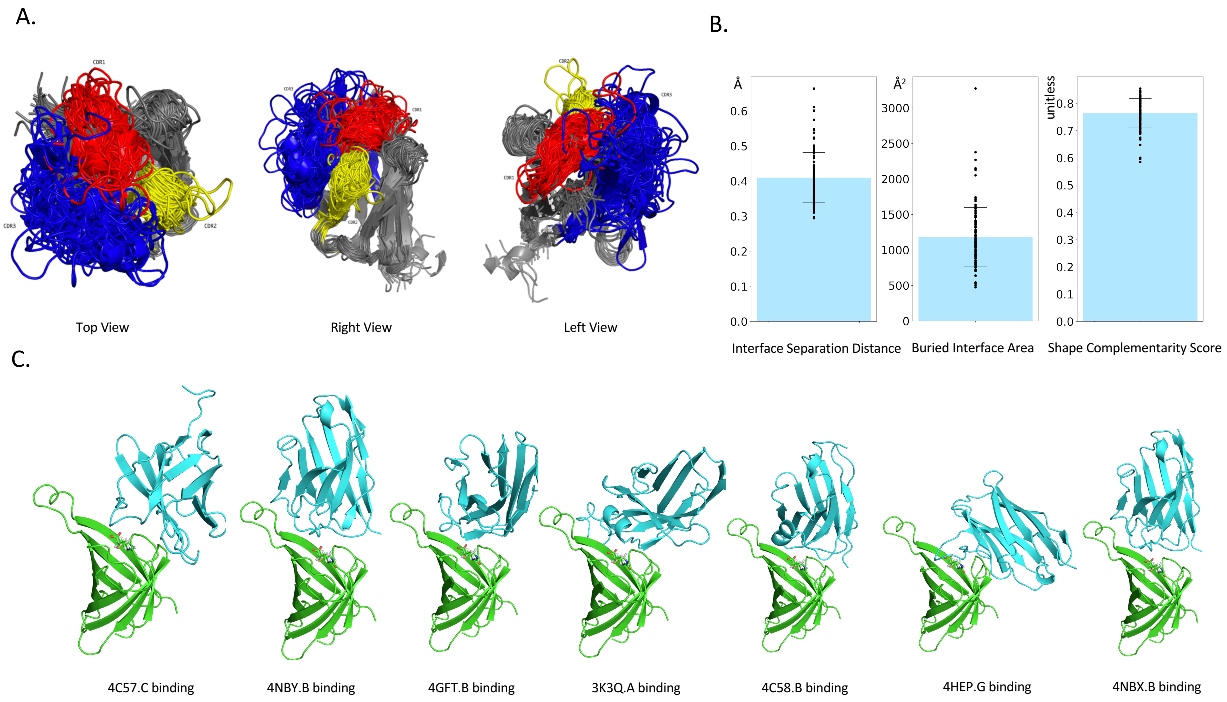


**Figure S1: Searching for optimal CDR conformation against monomeric streptavidin model.** (A). Whole structural-alignment of 154 curated PDB nanobody scaffolds with diverse CDR conformations and sequences. (B). Interface statistics of naturally occurring nanobody-target complexes. Error bars represent standard deviations. (C). The final 7 docked poses of nanobody scaffolds that passed the filters selecting poses that most likely recapitulate the natural binding modes of the corresponding nanobody scaffolds. Streptavidin S45A/T90A/D180A is colored green, and nanobody scaffold is colored cyan.

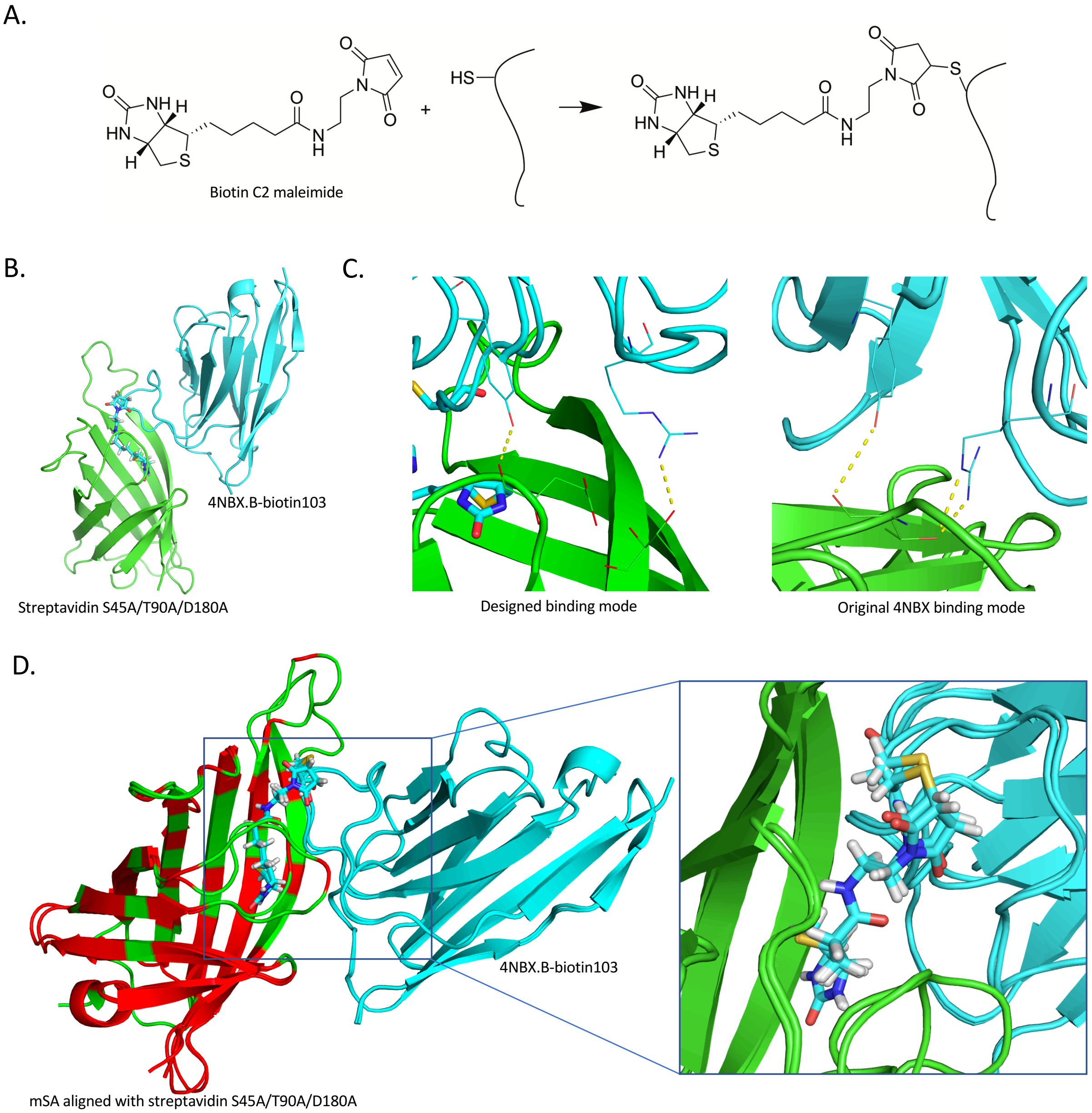


**Figure S2: Identification of optimal conjugation strategy and finalized conjugate models.** (A). Biotin conjugation was performed by biotin C2 maleimide with mutated cysteine residues. (B). Prepared structure of 4NBX.B-biotin103 in complex to streptavidin S45A/T90A/D180A. Streptavidin S45A/T90A/D180A is colored green, and nanobody scaffold is colored cyan. (C). The H-bond forming potential of Y112 and R27 in 4NBX.B nanobody was predicted to be recapitulated in the designed binding pose with the streptavidin model. Streptavidin S45A/T90A/D180A is colored green, and nanobody scaffold is colored cyan. Biotin103 side chain is shown as stick. Y112 and R27 together with their predicted H-bond partners are shown as line. (D). Alignment results for prepared structures of 4NBX.B-biotin103 in complex with streptavidin S45A/T90A/D180A and mSA. Streptavidin models are colored green, and nanobody scaffolds are colored cyan. Residues that were identified by sequence alignment as pair-wise identical sequences are colored red. Biotin103 side chain from both models are shown as stick.

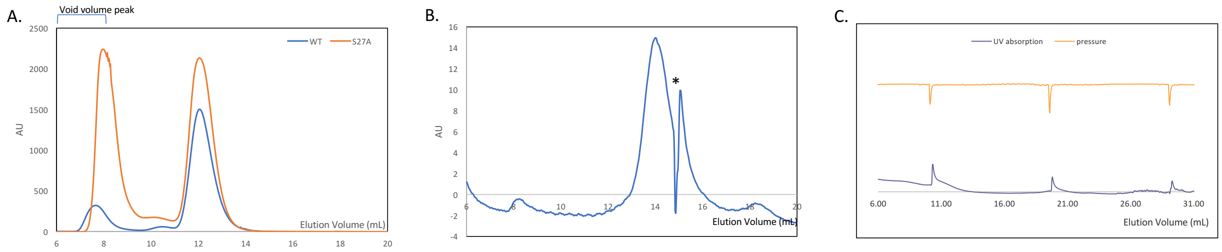


**Figure S3: Additional Supporting SEC traces.** A). SEC traces of $mSA_{WT}$ and $mSA_{S27A}$. B). SEC rerun trace of collected monomeric fraction for 4NBX.B-biotin103 v186_Fr. * indicates peaks of sample-irrelevant instrument defect of the overall FPLC, please refer to section C for more details. C). Blank run of the FPLC to reveal the sample-irrelevant periodic peaks that were constantly observed in SEC data.

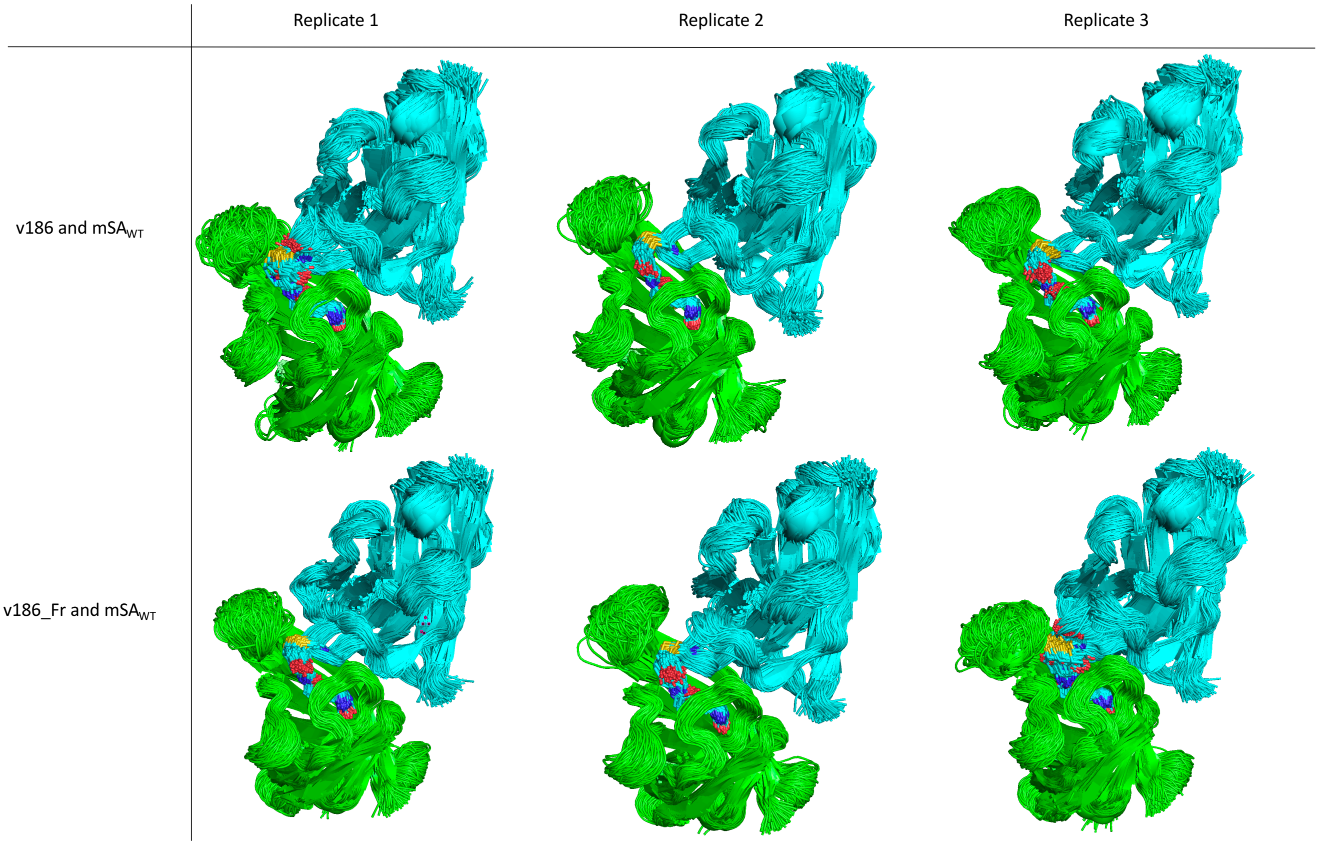


**Figure S4. Summary of MD simulations performed for 4NBX.B-biotin103 v186 and v186_Fr against mSA$_{WT}$.** For each simulation, 400 snapshots evenly spaced along the 100ns timescale are aligned together. green: mSA$_{WT}$. cyan: nanobody-biotin conjugates. The biotin103 "residue" is shown as stick.

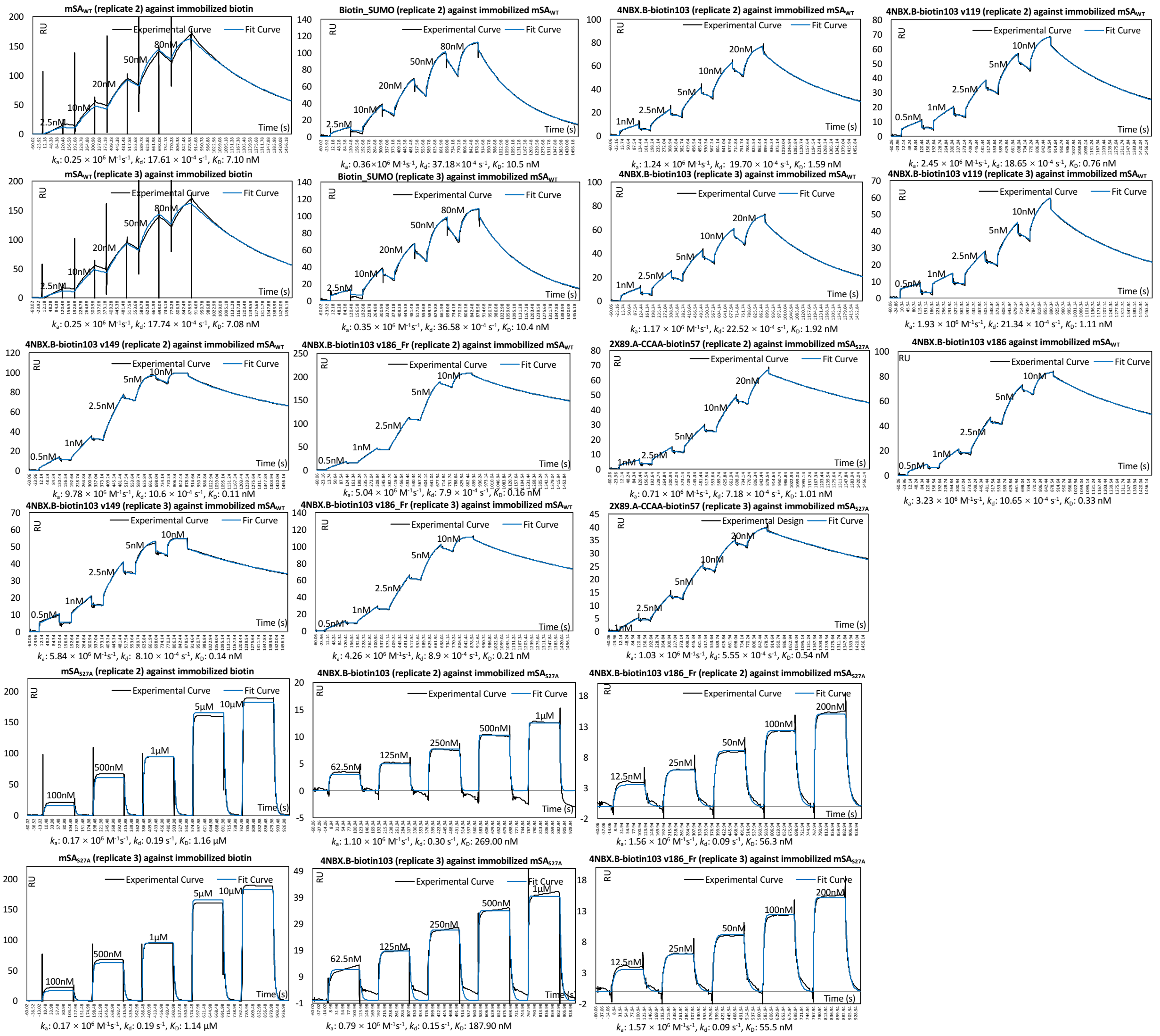


**Figure S5: SPR measurements from the intermediate design variant 4NBX.B-biotin103 v186, and from additional biological replicates not shown in the main text but were included for affinity and kinetics estimation.**

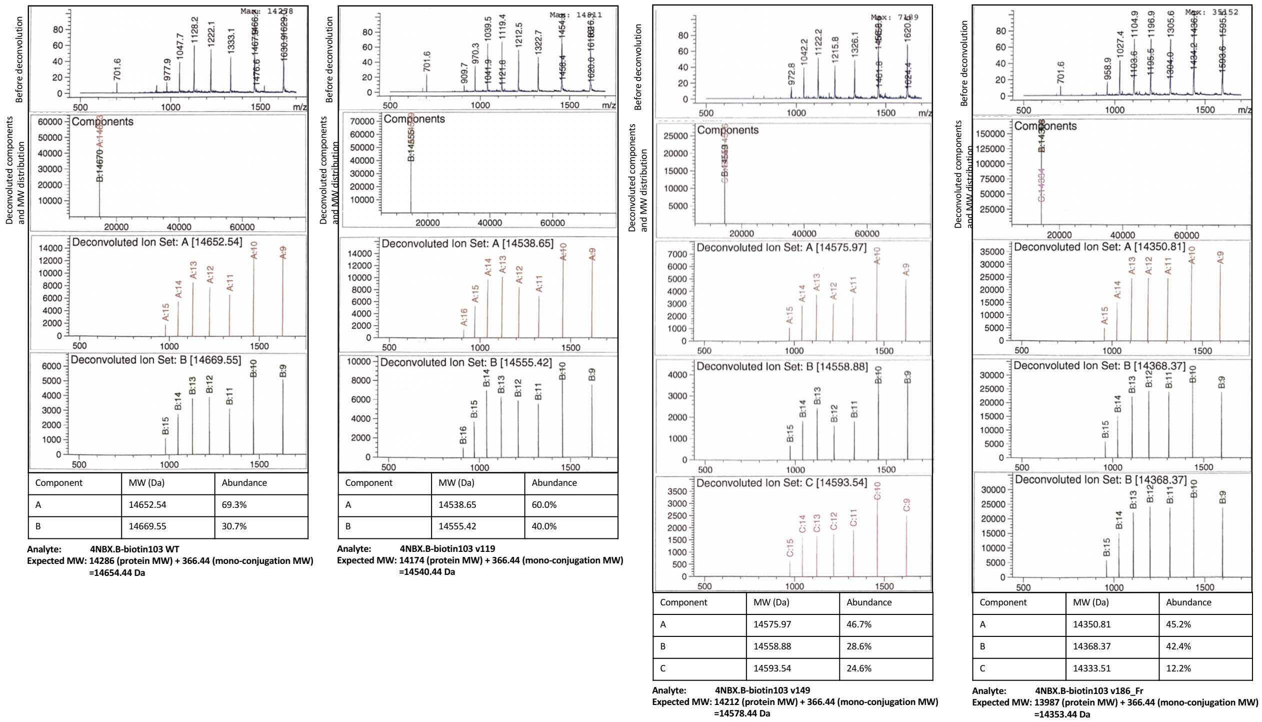


| Component | MW (Da) | Abundance |
|---|---|---|
| A | 14652.54 | 69.3% |
| B | 14669.55 | 30.7% |

| Component | MW (Da) | Abundance |
|---|---|---|
| A | 14538.65 | 60.0% |
| B | 14555.42 | 40.0% |

| Component | MW (Da) | Abundance |
|---|---|---|
| A | 14575.97 | 46.7% |
| B | 14558.88 | 28.6% |
| C | 14593.54 | 24.6% |

| Component | MW (Da) | Abundance |
|---|---|---|
| A | 14350.81 | 45.2% |
| B | 14368.37 | 42.4% |
| C | 14333.51 | 12.2% |

**Figure S6: Intact-protein mass spectrometry (MS) confirmed mono-conjugated materials.** MS deconvolution of nanobody-biotin conjugates only returned MWs within 20Da from expected MW of mono-conjugated materials.

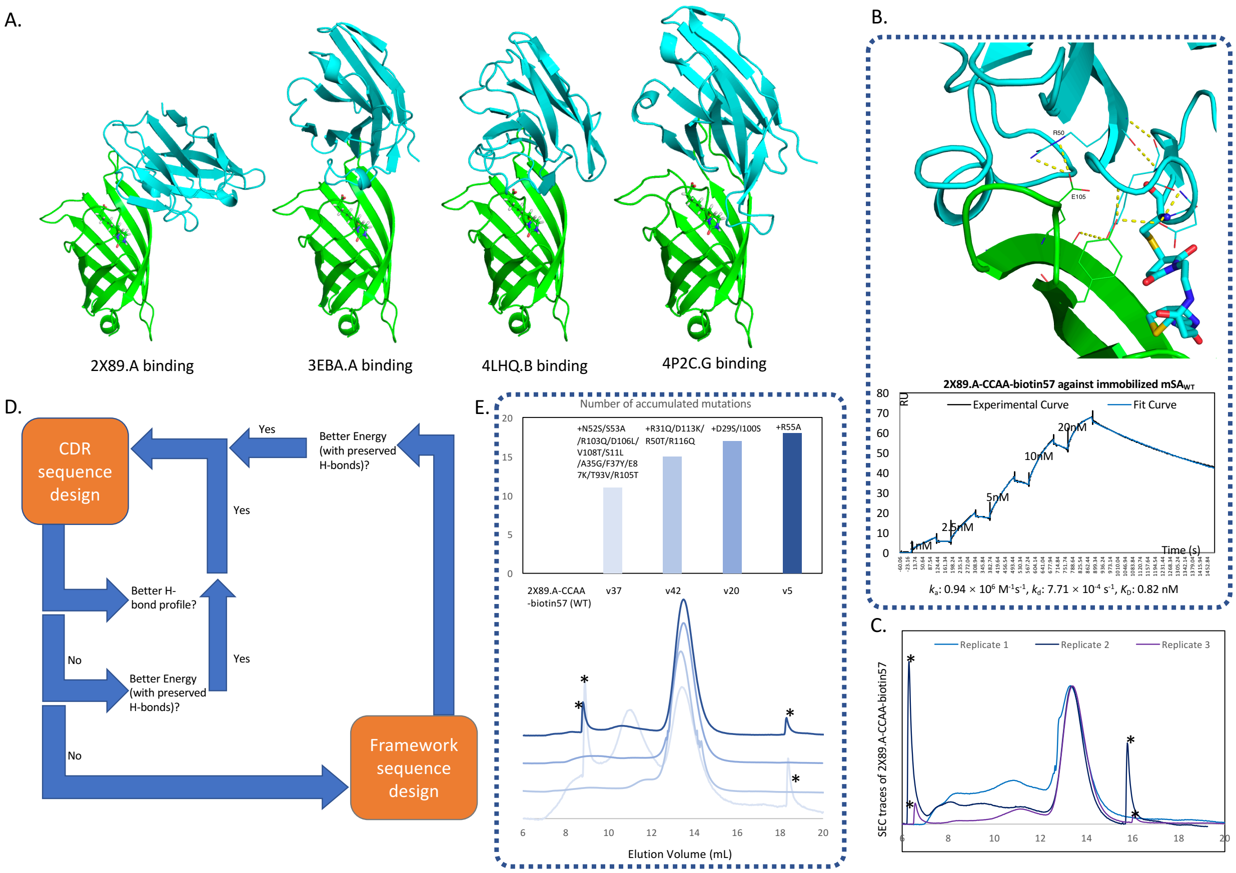


**Figure S7: Designing and testing of a nanobody scaffold obtained by directly docking against mSA.** (A). Finalized nanobody scaffolds and binding poses against mSA streptavidin. (B). Predicted H-bond formation profile and SPR binding curve for 2X89.A-CCAA-biotin57 WT against immobilized $mSA_{WT}$. Data from one of the triplicates is shown here, and data from the other two replicates is in Fig. S5. (C). Size-exclusion chromatography (SEC) traces of biological triplicates for 2X89.A-CCAA-biotin57, normalized by monomer peak height for better comparison of aggregates formation. * indicates peaks of sample-irrelevant instrument defect of the overall FPLC, please refer to figure S3C for more details. (D). A rudimentary sequence design pipeline that performs CDR and framework design in a step wise manner. (E). SEC traces and newly accumulated mutations for sequence-designed variants of 2X89.A-CCAA-biotin57 conjugates. Peak of the monomeric fractions were normalized to identical heights. * indicates peaks of sample-irrelevant instrument defect of the overall FPLC, please refer to figure S5C for more details.

**Table S1. Top 20 sequence outputs from CDR design of site 31, 32, 104, and 105 on 4NBX.B-biotin103 WT. (Ranked by energy score)**

| Ranking | Energy Score | Note | Mutations: chain name and accepted residue |
|---|---|---|---|
| 1 | -457.58628 | v119 | B_31H+B_32A+B_104S+B_105H |
| 2 | -456.82531 | | B_31N+B_32S+B_104S+B_105S |
| 3 | -456.56855 | | B_31N+B_32S+B_104S+B_105A |
| 4 | -456.56059 | | B_31R+B_32A+B_104D+B_105R |
| 5 | -456.50215 | | B_31R+B_32A+B_104S+B_105Y |
| 6 | -456.42944 | | B_31N+B_32A+B_104S+B_105S |
| 7 | -456.25964 | | B_31Q+B_32A+B_104S+B_105Y |
| 8 | -456.13421 | | B_31H+B_32S+B_104S+B_105V |
| 9 | -456.11428 | | B_31N+B_32A+B_104S+B_105R |
| 10 | -455.81205 | | B_31N+B_32A+B_104S+B_105A |
| 11 | -455.7737 | | B_31N+B_32S+B_104S+B_105D |
| 12 | -455.67248 | | B_31R+B_32A+B_104D+B_105H |
| 13 | -455.6018 | | B_31Q+B_32A+B_104A+B_105R |
| 14 | -455.26629 | | B_31R+B_104S+B_105H |
| 15 | -455.13137 | | B_31H+B_32S+B_104D+B_105S |
| 16 | -455.10018 | | B_31N+B_32S+B_104A+B_105S |
| 17 | -455.05911 | v149 | B_31R+B_32S+B_104A+B_105R |
| 18 | -454.97734 | | B_31R+B_32S+B_104S+B_105Y |
| 19 | -454.93878 | | B_31K+B_32S+B_104A+B_105A |
| 20 | -454.90643 | | B_31Q+B_32S+B_104A+B_105A |

Note: chain B refers to the nanobody

**Table S2. Top 20 sequence outputs from framework design on 4NBX.B-biotin103 v186. (Ranked by energy score)**

| Ranking | Energy Score | Note | Mutations: chain name and accepted residue |
|---|---|---|---|
| 1 | -500.8748 | v186_Fr | B_12V+B_37Y |
| 2 | -500.3636 | | B_37Y+B_40A |
| 3 | -500.24131 | | B_5Q+B_12V+B_37Y |
| 4 | -500.10634 | | B_12V+B_35G+B_37Y |
| 5 | -499.9846 | | B_12V+B_37Y+B_40A |
| 6 | -499.90705 | | B_12V |
| 7 | -499.22799 | | B_12V+B_35G |
| 8 | -499.15484 | | B_5Q+B_12V+B_35G+B_37Y |
| 9 | -499.02353 | | B_12V+B_35G+B_37Y+B_40A |
| 10 | -498.98295 | | B_5Q+B_12V+B_35G+B_37Y+B_40A |
| 11 | -498.78612 | | B_37Y |
| 12 | -498.6384 | | B_12V+B_35G+B_40A |
| 13 | -498.54874 | | B_35G+B_37Y |
| 14 | -498.53802 | | B_12V+B_40A |
| 15 | -498.48109 | | B_5Q+B_12V+B_40A |
| 16 | -498.31249 | | B_5Q+B_37Y |
| 17 | -498.04085 | | WT |
| 18 | -497.75764 | | B_5Q+B_12V |
| 19 | -497.68241 | | B_5Q |
| 20 | -497.59715 | | B_5Q+B_35G+B_37Y |

Note: chain B refers to the nanobody

**Table S3. Top 20 sequence outputs from framework design on 4NBX.B-biotin103 v149. (Ranked by energy score)**

| Ranking | Energy Score | Note | Mutations: chain name and accepted residue |
|---|---|---|---|
| 1 | -482.29049 | | WT |
| 2 | -481.67807 | | B_37Y |
| 3 | -481.6627 | | B_5Q+B_12V |
| 4 | -481.65469 | A12V/F37Y mutations | B_12V+B_37Y |
| 5 | -481.51508 | | B_12V+B_35G |
| 6 | -481.5122 | | B_5Q |
| 7 | -481.48205 | | B_35G |
| 8 | -481.38994 | | B_35G+B_37Y |
| 9 | -481.32699 | | B_5Q+B_37Y |
| 10 | -481.13866 | | B_40A |
| 11 | -481.12555 | | B_12V |
| 12 | -480.83661 | | B_35G+B_40A |
| 13 | -480.82825 | | B_12V+B_37Y+B_40A |
| 14 | -480.50684 | | B_5Q+B_35G+B_37Y |
| 15 | -480.44015 | | B_5Q+B_12V+B_40A |
| 16 | -480.42843 | | B_12V+B_35G+B_37Y |
| 17 | -480.3031 | | B_5Q+B_12V+B_37Y+B_40A |
| 18 | -480.19273 | | B_5Q+B_12V+B_35G+B_40A |
| 19 | -480.16999 | | B_5Q+B_35G+B_40A |
| 20 | -480.16866 | | B_12V+B_35G+B_37Y+B_40A |

Note: chain B refers to the nanobody

**Table S4. amino acid sequences for the protein templates used in this study**

| | |
|---|---|
| 4NBX.B_A103C | QVQLQESGGGLAQAGGSLRLSCAASGRTFSMDPMAWFRQPPGKEREFVAAGSSTGRTTYYADSVKGRFTISRDNAKNTVYLQMNSLKPEDTAVYYCAAAPYGCNWYRDEYAYWGQGTQVTVSSHHHHHH |
| 2X89.A_CCAA_I57C | QVQLQESGGGSVQAGGSLRLSCAASGYTDSRYAMAWFRQAPGKEREWVARINSGRDCTYYADSVKGRFTFSQDNAKNTVYLQMDSLEPEDTATYYCATDIPLRARDIVAKGGDGFRYWGQGTQVTVSSHHHHHH |
| mSA_WT | HHHHHHSQDLASAEAGITGTWYNQSGSTFTVTAGADGNLTGQYENRAQGTGCQNSPYTLTGRYNGTKLEWRVEWNNSTENCHSRTEWRGQYQGGAEARINTQWNLTYEGGSGPATEQGQDTFTKVKPSAASGSDYKDDDDK |
| Smt3 SUMO with N-terminal cysteine | CLQDSEVNQEAKPEVKPEVKPETHINLKVSDGSSEIFFKIKKTTPLRRLMEAFAKRQGKEMDSLRFLYDGIRIQADQAPEDLDMEDNDIIEAHREQIGGHHHHHH |

Note: the numbering scheme of the above sequences follows 1-2-3-4… The first residue is numbered as 1.
